# Differential regulation of mouse hippocampal gene expression sex differences by chromosomal content and gonadal sex

**DOI:** 10.1101/2021.09.01.458115

**Authors:** Sarah R. Ocañas, Victor A. Ansere, Kyla B. Tooley, Niran Hadad, Ana J. Chucair-Elliott, David R. Stanford, Shannon Rice, Benjamin Wronowski, Kevin D. Pham, Jessica M. Hoffman, Steven N. Austad, Michael B. Stout, Willard M. Freeman

## Abstract

Common neurological disorders, like Alzheimer’s disease (AD), multiple sclerosis (MS), and autism, display profound sex differences in prevalence and clinical presentation. However, sex differences in the brain with health and disease are often overlooked in experimental models. Sex effects originate, either directly or indirectly, from hormonal or sex chromosomal mechanisms. To delineate the contributions of genetic sex (XX v. XY) versus gonadal sex (ovaries v. testes) to the epigenomic regulation of hippocampal sex differences, we use the Four Core Genotypes (FCG) mouse model to uncouple chromosomal and gonadal sex. Transcriptomic and epigenomic analyses of ∼12-month-old FCG mice hippocampi, revealed genomic context-specific regulatory effects of genotypic and gonadal sex on X- and autosome-encoded gene expression and DNA modification patterns. X-chromosomal epigenomic patterns, classically associated with X-inactivation, were established almost entirely by genotypic sex, independent of gonadal sex. Differences in X-chromosome methylation were primarily localized to gene regulatory regions including promoters, CpG islands, CTCF binding sites, and active/poised chromatin, with an inverse relationship between methylation and gene expression. Whereas, autosomal gene expression demonstrated regulation by both genotypic and gonadal sex, particularly in immune processes. These data demonstrate an important regulatory role of sex chromosomes, independent of gonadal sex, on sex-biased hippocampal transcriptomic and epigenomic profiles. Future studies will need to further interrogate specific CNS cell types, identify the mechanisms by which sex chromosomes regulate autosomes, and differentiate organizational from activational hormonal effects.

## Introduction

Sex is a major risk factor for many neurological diseases and disorders, including Alzheimer’s disease (AD) [1, 2], multiple sclerosis [3, 4], autism [5], attention-deficit/hyperactivity disorder (ADHD) [6], depression [7], and age-related cognitive decline [8, 9]. Of particular importance in modulating the cognitive effects seen in many sex-biased diseases is the hippocampus, the learning center of the brain [10]. Females tend to outperform males on hippocampal-dependent learning tasks and are more impacted by many diseases/disorders of hippocampal dysfunction (i.e. AD, depression) [11]. Understanding basal sex differences and their regulation in the hippocampus can help gain insight into the etiology of sex differences in hippocampal dysfunction in common neurological diseases. The goal of this study is to separate the effects of gonadal and chromosomal sex on the sex differential regulation of mouse hippocampal gene expression through epigenomic mechanisms.

Sex characteristics can be influenced by genotypic sex, gonadal sex, and gender. Generally, in mammals, a genotypic female has two X-chromosomes (and no Y-chromosome), while a genotypic male has one X-chromosome and one Y-chromosome. Gonadal sex is classified based on the individual’s genitalia, which can influence expression of secondary sex characteristics [12]. Gonadal sex determination is driven by the presence or absence of the Y-chromosome encoded sex-determining region of Y (*Sry*) gene. *Sry* is necessary and sufficient for development of testes, and in the absence of *Sry* mammals develop ovaries [13]. As such, genotypic and gonadal sex are causally linked and the relative contributions of chromosomal and gonadal sex to sex-biases in health and disease are difficult to disentangle.

On the other hand, gender is a societal construct that can be molded by an individual’s perception of their sex, as well as influences from their social and physical environments. Although gender, including behavioral and societal influences, likely impacts health and disease outcomes [14], it is not possible to discern gender in non-human animal models. Here we focus on the relative contributions of genotypic and gonadal sex to sex differences in the adult mouse hippocampus.

After gonad differentiation, hormonal secretions influence the organism’s sexual phenotype. Gonadal hormonal secretions lead to organizational effects that cause sex differentiation during development, as well as activational effects that may be temporary and reversible and can occur at any stage of life [15]. In a seminal paper, Phoenix et al (1959) [16] described the organizational-activational theory of sexual differentiation in which during early development, hormones have an organizational effect on neural tissue development and circuitry that mediates mating (and likely other behaviors). After the organizational framework is established during development, activational effects are mediated by levels of gonadal and non-gonadal sex hormones. As such, the direct contributors to phenotypic sex effects are: (1) activational effects of gonadal hormones, (2) organizational effects of gonadal hormones, and (3) sex chromosomal effects [17].

Since the onset of developmental and age-related brain diseases correlate to critical windows of hormonal transition, gonadal hormone effects have been widely studied in brain diseases [18–22]. However, inconsistent efficacy of hormonal replacement therapies (HRT) in diseases like AD [23–25], potential adverse outcomes [26, 27], and the feminizing/masculinizing effects of gonadal hormones have diminished enthusiasm for HRT approaches. On the other hand, several studies have also pointed to distinct contributions of sex chromosomes to brain development [28, 29] and pathology [30–32]. Thus, establishing the contributions of sex chromosomes, independent of gonadal hormones, to brain diseases have come to the forefront of the neurobiology of sex differences.

Despite being the largest genomic difference between humans, the role of sex chromosomes in regulating sex effects is complex and still mostly unclear. Although the human Y-chromosome only contains 568 genes (71 protein-coding) [33], mosaic loss of Y with age has been linked to cancer [34] as well as AD [35]. In comparison, the X-chromosome has between 900 and 1500 genes, but its expression profile is complicated by the random inactivation in females of one X-chromosome on a cell-by-cell basis due to dosage compensation [36]. In fact, the complex nature of sex chromosome genomic regulation led to standardized exclusion of sex chromosomes from genome wide association studies (GWAS) [37], even those specifically interested in sex effects with brain disease [38]. X-chromosome gene dosage effects are evident in human sex chromosome aneuploidies, including Turner syndrome (XO), Triple X Syndrome (XXX), and Klinefelter syndrome (XXY), each with a wide array of CNS symptomology [39–41]. Thus, in humans it appears that genotypic sex, especially the number of X-chromosomes, plays an important role in proper brain development and function. However, sex chromosomal aneuploidies in humans are often confounded by different levels of gonadal hormones. Whereas, in mouse models the levels of hormones are more easily controlled, and can be dissociated from the effects of different numbers of sex chromosomes.

To disentangle the effects of gonadal sex (testes v. ovaries; M v. F) and sex chromosome complement (XX v. XY) on the steady state gene expression and DNA modification patterning of the hippocampus, we use the Four Core Genotypes (FCG) mouse model [42]. The FCG male (XYM) originated through two sequential genetic changes [43]: 1) a spontaneous deletion of the testis-determining *Sry* gene from the Y-chromosome (Y^-^*^Sry^*) [44] and 2) transgenic insertion of the *Sry* gene onto an autosome (A*^Sry^*) [45] resulting in XY^-^*^Sry^*A*^Sry^* (XYM). Crossing the XYM with a wild-type C57BL/6J XX female (XXF), results in the uncoupling of gonadal and chromosomal sex: XX and XY^-^*^Sry^* mice with ovaries (XXF/XYF) and XXA^Sry^ and XY^-^*^Sry^*A^Sry^ mice with testes (XXM/XYM) (**Fig. 1A**). This allows for 2-way statistical comparisons to assess the contributions of gonadal and chromosomal sex, as well as interactive effects, on molecular and phenotypic outcomes (**Fig. 1B**).

**Fig. 1.**
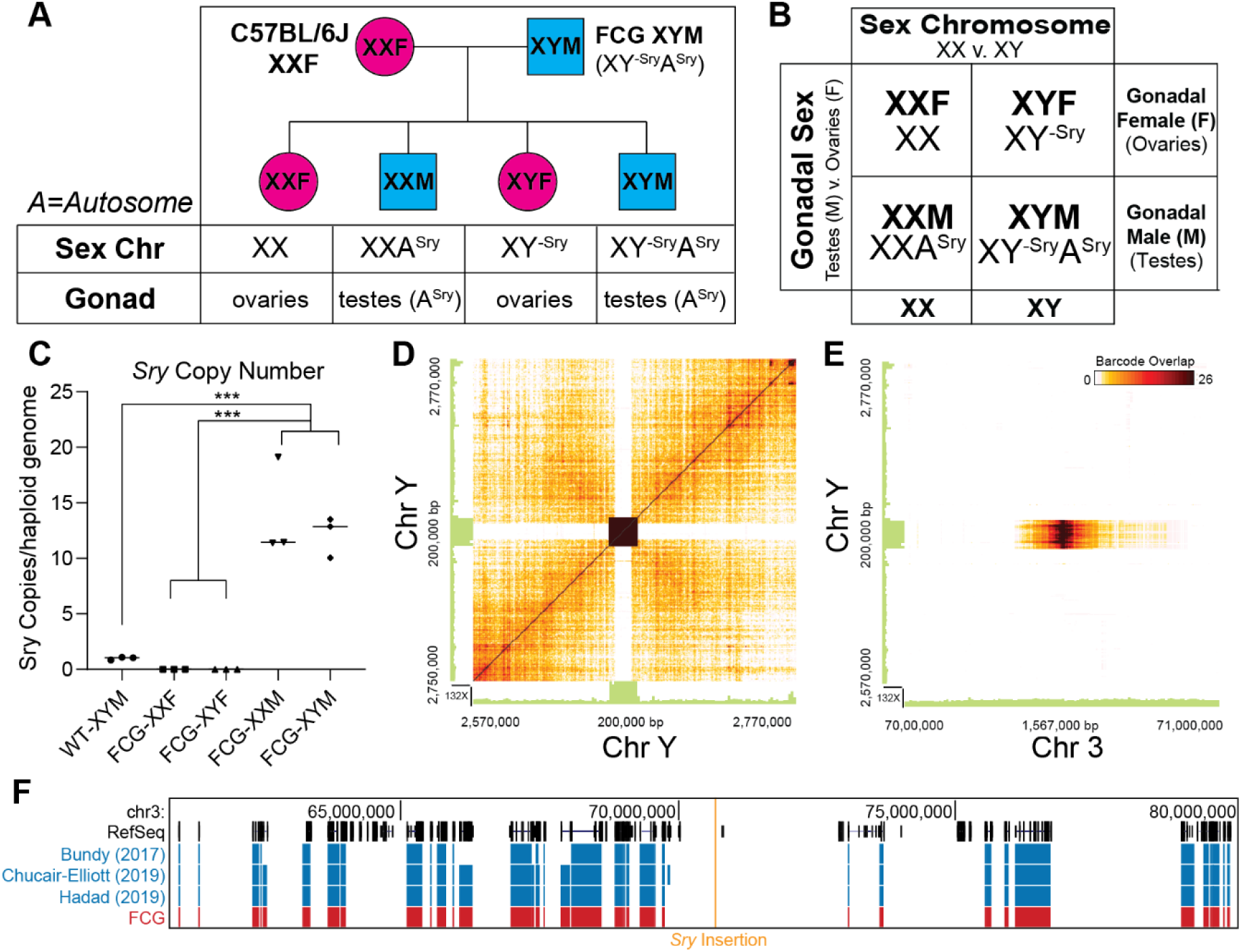
*Sry* copy number and localization in FCG hippocampi. **A**) Breeding strategy used to generate the Four Core Genotypes (FCG). **B**) FCG mouse model two-way design to study gonadal sex and sex chromosomal contributions to sex effects. DNA isolated from FCG and wild type hippocampi (n=3/group) was used for digital PCR *Sry* copy number and 10X genomics linked read sequencing. **C**) Wildtype (WT) C57BL/6J mice have one copy of the testis-determining *Sry* gene, while FCG males (both XX and XY) have 12-14 copies of *Sry. Sry* was not detected in the FCG females. **D**) Linked read sequencing of FCG XYM, shows strong linkage of the *Sry* gene to itself, but not to adjacent regions of the Y-chromosome (ChrY) indicating *Sry* is no longer on ChrY. **E**) *Sry* gene shows strong linkage to an intergenic region on Chromosome 3 (Chr3). **F)** Adult mouse (3-4 months old) hippocampal gene expression from previously published studies [Bundy et al., 2017 (GSE83931), Chucair-Elliott et al., 2019 (GSE135752); and Hadad et al., 2019 (PRJNA523985)] were compared to FCG hippocampal gene expression in the present study, to determine potential alterations in gene expression adjacent (+/- 10 Mbp) to the *Sry* insertion site on Chr3 in the FCG XXM/XYM. Tracks represent genes that are detected as expressed in previously published datasets (blue) and the present FCG study (red).

Early FCG mouse studies were aimed at determining if sex chromosome complement (XX v. XY) contributed to development of well-established sexually dimorphic phenotypes [43, 46–48] [49] [50, 51]. Later FCG studies have brought to light distinct contributions of genotypic sex to disease-associated phenotypes seen in Experimental Autoimmune Encephalomyelitis (EAE) and pristane-induced Systemic Lupus Erythematosus (SLE) mouse models [30]. Additionally, XX mice have longer lifespans than their XY counterparts, regardless of gonadal sex [31], and XX mice show resiliency to death in an AD mouse model [32].

Although hippocampal sex differences in the transcriptome and epigenome are well-established across development, aging, and disease in mice and humans, the relative contributions of sex chromosome complement (XX v. XY) and gonadal sex (M v. F) to the steady state and stimulus responsive transcriptome and epigenome are not fully defined. Given its influence on X-chromosome inactivation (XCI), genome accessibility, and regulation of gene expression, specific methylation of the fifth carbon of a cytosine residue, resulting in the DNA modification 5-methyl-cytosine (mC) is a particularly interesting epigenetic factor. In this study, we use transcriptomic and epigenetic approaches to examine the hippocampal transcriptome and methylome in adult FCG mice. We then compare our findings to previously identified hippocampal sex differences to begin to separate contributions of sex chromosome complement (XX v. XY) and gonadal sex (testes v. ovaries; M v. F) to sex differences in transcriptional programming.

## Results

### *Sry* copy number and localization in adult FCG hippocampi

The testis-determining *Sry* gene is considered the “master switch” in mammalian gonadal sex determination [52]. In the FCG model, *Sry* is absent from the Y-chromosome and inserted onto an autosome, uncoupling gonadal and genetic sex. Previous initial reports [53] used fluorescence *in situ* hybridization (FISH) and PCR amplicon sequencing to localize a concatemer (12-14 copies) of *Sry* on chromosome 3 (Chr3: 70673749-70673824) in FCG XX and XY males. To verify *Sry* copy number, we designed a digital PCR *Sry* copy number assay and confirmed 12-14 copies of *Sry* in FCG males (XXM/XYM), as compared to one copy in WT males (**Fig. 1C**). To confirm the localization of *Sry,* a form of linked-read sequencing was used, which barcodes sequence reads that come from the long DNA fragments with the same oligo tag. Within the visualization, a darker amber color indicates more linked reads between the regions on the X and Y axes (**Fig. 1D**). There was no linkage between *Sry* and the adjacent region of the Y-chromosome, indicating deletion of *Sry* from the Y-chromosome. Strong linkage of the *Sry* gene with the previously-identified [53] region of chromosome 3 (**Fig. 1E**) within a region with no known gene annotation. No other linked regions were identified indicating that this is the only autosomal insertion site of *Sry*. To examine if *Sry* insertion alters expression of genes on Chr3, hippocampal RNA-Seq data from and wild-type C57Bl/6J mice was compared for genes adjacent (+/- 10Mbp) to the *Sry* insertion site **(Fig. 1F)**. Forty genes were expressed at detectable levels (>20 reads) in male and female hippocampi in all three previous sex difference studies examined [54–56], and each of these 40 genes were also expressed across all four groups in the FCG hippocampi (**Supplemental Table 1**). No sex differences were identified across the three previously published studies. Only a single gene (*Fam198b*) ∼9.2 Mbp from the *Sry* insertion was differentially expressed by sex in the FCG hippocampi. Due to the distance from the insertion site and variability in autosomal sex differences, this sex difference is unlikely a result of the *Sry* insertion. Further, there was no ectopic expression of *Sry* in the hippocampi of male FCG mice.

### Transcriptomic analysis of sex chromosomal (X/Y) differential expression from adult FCG hippocampi

Despite X-inactivation compensatory mechanisms, there are a number of X-chromosome genes whose expression is imbalanced between males and females in the mouse hippocampus. Previous reports have established differentially expressed sex chromosomally-encoded genes in the mouse hippocampus throughout development and aging [54–57] (**Supplemental Table 2; GEO Accession:** GSE83931, GSE135752, GSE76567**; SRA bioProject:** PRJNA523985). Intersecting sex chromosomally-encoded (X/Y) differentially expressed genes by sex across studies identified eight common genes (**Fig. 2A, Supplemental Table 2**), including X-chromosome genes (*Xist, Ddx3x, Kdm6a,* and *Eif2s3x*) and Y-chromosome genes (*Kdm5d, Eif2s3y, Uty, Ddx3y*). The four common ChrX genes (*Xist, Ddx3x, Kdm6a,* and *Eif2s3x*) have all been identified as genes likely to escape XCI [58]. XCI escape of *Ddx3x*, *Kdm6a*, and *Eif2s3x* are likely due to intolerance of haploinsufficiency, as each of these genes have Y-encoded paralogs *Ddx3y, Uty, and Eif2s3y*, respectively.

**Fig. 2.**
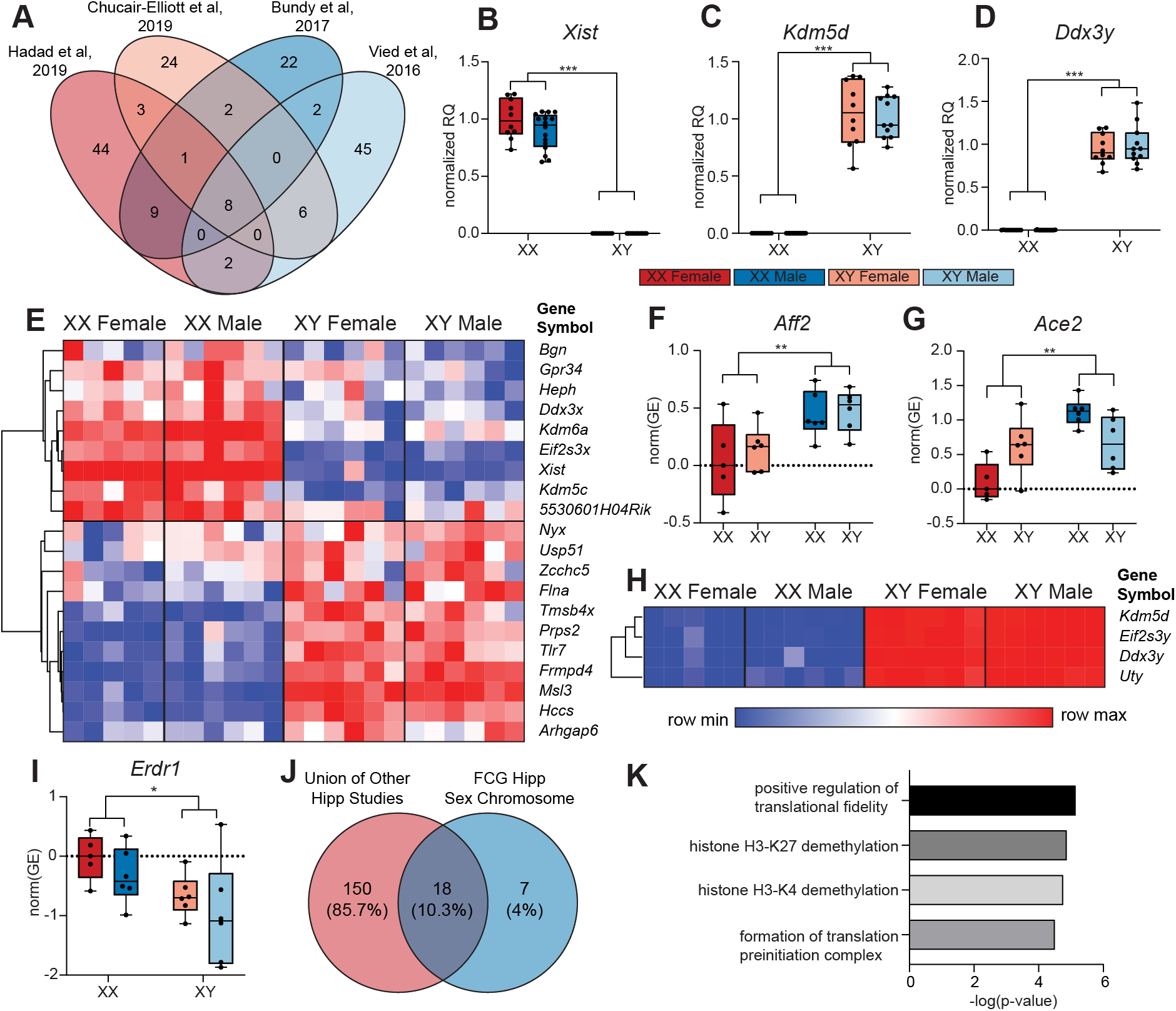
Transcriptomic analysis of differential expression in sex chromosome-encoded (X/Y) genes in adult FCG hippocampi. DNA and RNA were isolated from FCG hippocampi (n=10-16/group). mRNA expression was assessed by RT-qPCR (n=10-16/group) and stranded RNA-Seq (n=6/group). Results were compared to previously published hippocampal transcriptomic sex differences. **Boxplots** represent median, interquartile range, and minimum/maximum normalized gene expression. **A**) Comparison of four previous hippocampal transcriptomic studies, showed 168 sex chromosome-encoded sex differences in wild-type mice across studies with eight genes common between all studies (*Xist Ddx3x, Kdm6a, Eif2s3x, Kdm5d, Eif2s3y, Uty, Ddx3y)* **B**) In the FCG hippocampus, RT-qPCR of X-chromosome encoded gene *Xist* confirms differential expression by sex chromosome (XX vs. XY) but not gonadal sex (M v. F) (Two-way ANOVA, main effect of sex-chromosome complement (XX v. XY), ***p<0.001) . **C-D**) RT-qPCR of Y-chromosome encoded genes shows similar levels of expression of (C) *Kdm5d* and (D) *Ddx3y* in XYM and XYF, and no detectable expression in XXM or XXF (Two-way ANOVA, main effect of sex chromosome complement (XX v. XY), ***p<0.001). **E**) RNA-Seq analysis of X-chromosome encoded genes showed 20 genes that are differentially expressed by sex chromosome (XX vs. XY) but not by gonadal sex (M v. F). **F-G**) RNA-seq analysis of X chromosome genes revealed only two genes (*Aff2, Ace2*) differentially expressed by gonadal sex (M v. F) and not by sex chromosome complement. Both (**F**) *Aff2* and (**G**) *Ace2* had higher expression in gonadal males than females regardless of their sex chromosome (Two-way ANOVA, main effect of gonadal sex (M v. F), **p<0.01). **H-I**) RNA-Seq analysis of Y-chromosome encoded genes identified 5 differentially expressed genes by sex chromosome (XX vs. XY) but not by gonadal sex (M v. F). There were no Y-chromosome genes that were differentially expressed by gonadal sex (M v. F). **H**) Four of the genes (*Kdm5d, Eif2s3y, Ddx3y, Uty*) show no expression in XX genotypes. **I**) Located in the pseudo autosomal region (PAR) of the X/Y-chromosomes, *Erdr1* shows higher expression in XX genotype. **J**) Comparing the union of previous hippocampal studies described in (**A**) to the FCG sex chromosome genes differentially expressed between XX and XY genotypes, yields 18 common genes. **K**) GO Ontology analysis of the 18 genes from (**J**), identified four significantly enriched biological pathways

Common sexually dimorphic genes from these previous studies were examined in FCG (XXF, XYF, XXM, XYM) hippocampi (n=10-16/group) by RT-qPCR to examine gonadal versus chromosomal sex regulation. X-inactive specific transcript (*Xist*), a non-coding RNA, stabilizes the inactive X-chromosome (Xi) and interacts with various silencing factors to alter chromatin accessibility through a variety of epigenetic mechanisms [59]. In the FCG hippocampus, *Xist* was differentially expressed by sex chromosome complement (XX v. XY) regardless of gonadal sex, with no detected expression in XY genotypes (**Fig. 2B**; Two-way ANOVA, Main effect of sex chromosome complement (XX v. XY), ***p<0.001) and no effect of gonadal sex or interactive effect of gonadal and chromosomal sex showing a clear influence of sex chromosome complement in the modulation of X-chromosome dosage compensation.

Y-encoded lysine demethylase 5D (*Kdm5d*) mediates H3K4 demethylation and regulates sexually dimorphic gene expression [60]. DEAD-Box Helicase 3 Y-Linked (*Ddx3y*) is part of the male-specific region of the Y-chromosome and contains a conserved Asp-Glu-Ala-Asp (DEAD) motif that is used by ATP-dependent RNA helicases [61]. Expression of Y-chromosome encoded genes *Kdm5d* and *Ddx3y* were assessed by RT-qPCR. In the FCG hippocampus, there was a main effect of chromosome (XX v. XY) on *Kdm5d* and *Ddx3y* expression, with higher expression in XY and no detected expression in XX genotypes (**Fig. 2C-D**; Two-way ANOVA, main effect of sex chromosome complement (XX v. XY), ***p<0.001). There was no main effect of gonadal sex (M v. F) or interactive effect of gonadal and chromosomal sex on the expression of *Kdm5d* or *Ddx3y* in the FCG hippocampus.

To assess the transcriptome in an unbiased manner, directional RNA Sequencing (RNA-Seq) libraries were prepared from FCG hippocampal RNA (n=5-6/group). After aligning, quantifying, and calling differentially expressed genes, twenty differentially expressed X-chromosome genes were identified with a main effect of sex chromosome (XX v. XY) regardless of gonadal sex (M v. F) with nine genes higher and 11 genes lower in XX vs. XY (**Fig. 2E, Supplemental Table 3**; Two-Way ANOVA, BHMTC, FC>=1.25, FDR<0.1). Only two X-chromosome genes (*Ace2, Aff2*) were differentially expressed by gonadal sex (M v. F) (**Fig. 2F-G, Supplemental Table 3**; Two-Way ANOVA, BHMTC, FC>=1.25, FDR<0.1). Four Y-encoded genes were found to be expressed and each of the four Y-chromosome genes were differentially expressed by chromosome (XX v. XY) (**Fig. 2H, Supplemental Table 3**; Two-Way ANOVA, BHMTC, FC>=1.25, FDR<0.1). There were no Y-chromosome genes differentially expressed by gonadal sex (M v. F). There were no X- or Y-chromosomal genes identified with interactive effects of gonadal and chromosomal sex in the FCG hippocampi.

Erythroid differentiation regulator 1 (*Erdr1*) is part of X- and Y-chromosome pseudoautosomal region (PAR) that is able to crossover and recombine during meiosis [62]. As such, genes within the PAR have the same sequence on the X- and Y-chromosome and the chromosomal origin (X or Y) of these transcripts cannot be determined with traditional RNA-Seq. In FCG hippocampi, *Erdr1* is differentially expressed by sex chromosome complement (XX v. XY), with higher levels in XX animals (**Fig. 2I**).

The eight common previously identified X/Y-chromosome sex differences (**Fig. 2A, Supplemental Table 2 (Intersection)**) were all differentially expressed by sex chromosome complement (XX v. XY) but not gonadal sex in the adult FCG hippocampus. When comparing the union of all sex differentially expressed genes from previous hippocampal studies (**Fig. 2A, Supplemental Table 2 (Union)**) to the differentially expressed genes by sex chromosome complement (XX vs. XY) in the FCG hippocampus (**Fig. 2B-I, Supplemental Table 3**) there were 18 genes in common (**Fig. 2J**). Functional analysis of differentially expressed genes by sex across all studies revealed enrichment in four overrepresented biological processes (Fisher’s Exact, FDR<0.05), including: 1) positive regulation of translational fidelity, 2) histone H3-K27 demethylation, 3) histone H3-K4 demethylation, and 4) formation of translation preinitiation complex (**Fig. 2K**). Regulation of histone H3-K27 and H3-K4 methylation have both been implicated in the initiation and maintenance of XCI [63].

Together, these data suggest that sex chromosome gene expression is tightly regulated by the sex chromosomes themselves, independent of gonadal sex. Further, sex chromosomally-encoded histone modifiers are likely involved in the maintenance of XCI and also may alter the autosomal epigenome.

### Sex chromosome (X/Y) methylation patterns by Whole Genome Oxidative Bisulfite Sequencing (WGoxBS) in FCG hippocampi

DNA methylation can regulate gene expression by a variety of mechanisms, including (but not limited to): 1) direct transcription inhibition by blocking transcription factor binding [64], 2) indirect transcription regulation by recruitment of chromatin modifiers and methyl binding proteins [65–68], 3) genomic imprinting [69], and 4) XCI [70–72]. Although methylated DNA is generally associated with transcriptional silencing, there are reported cases where DNA methylation may also serve an activational role [73].

In females, XCI occurs through multi-layer epigenetic mechanisms that ultimately compact the inactive X-chromosome (Xi) into a heterochromatic Barr Body. Early in development, the long-noncoding RNA (lncRNA) *Xist* is expressed from Xi and provides a *cis*-coating that recruits protein complexes, leading to changes in chromatin accessibility and DNA modifications [36]. Changes in histone modifications [63] and DNA methylation stabilizes Xi in the inactive state [74]. In this study, we assessed the efficacy of Xi epigenetic silencing in the FCG mouse model by analyzing the X chromosome DNA methylation patterning in FCG hippocampal DNA. DNA isolated from FCG hippocampi (n=3/group) was oxidized and bisulfite-converted prior to constructing whole genome libraries for sequencing. Oxidative bisulfite sequencing provides a specific methylation quantitation, avoiding comingling of hydroxymethylation signal. After aligning and calling methylation values, the whole genome methylation levels in both CG and non-CG (CH) contexts were calculated. There were no observed differences in overall whole genome (including autosomes and sex chromosomes) methylation in CG context (mCG) (**Fig. 3A**) by gonadal sex (M v. F) or sex chromosome complement (XX v. XY), consistent examinations of sex differences in mCG in WT mouse hippocampi [55].

**Fig. 3.**
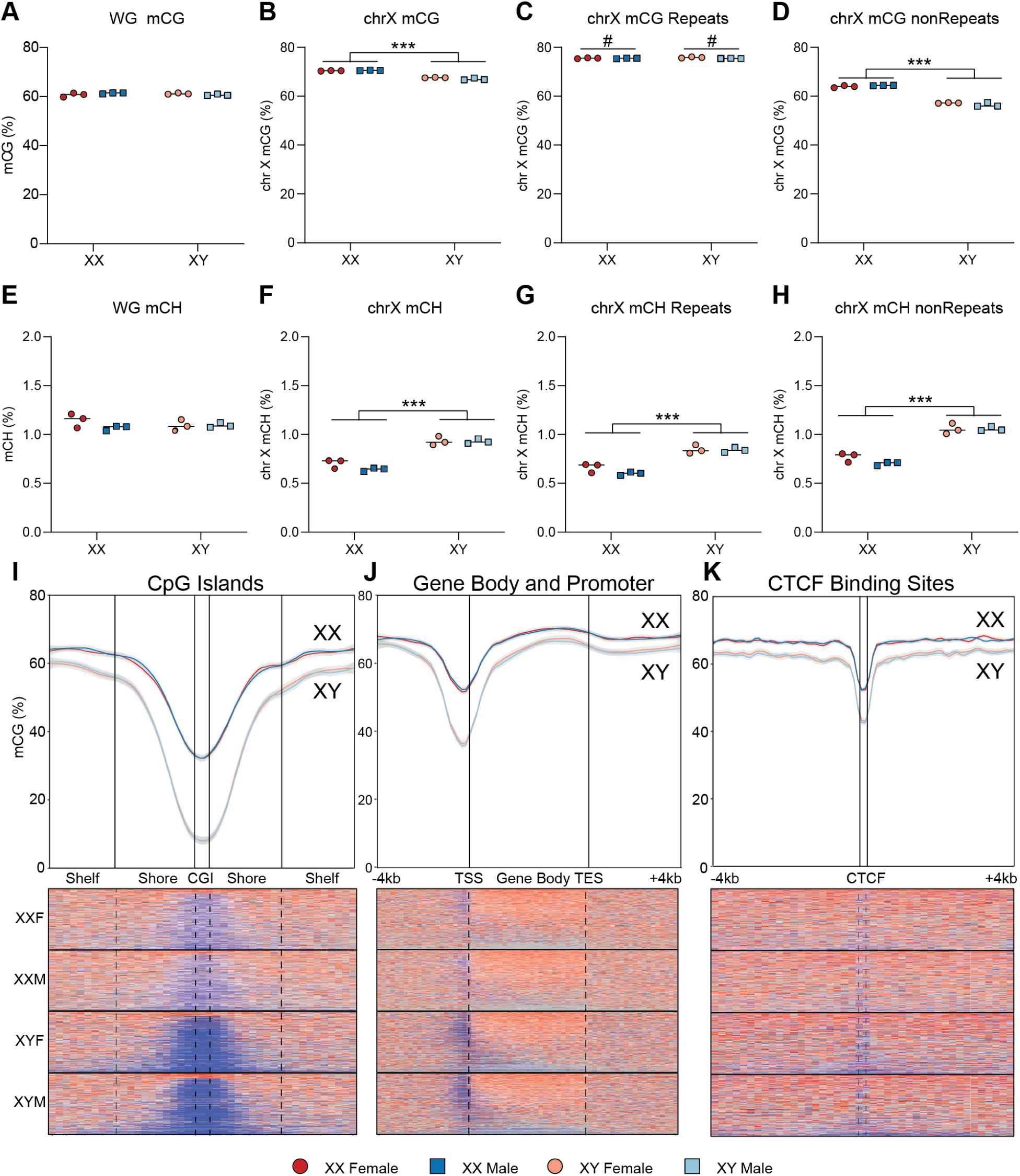
X-chromosome DNA methylation landscape in FCG hippocampus by WGoxBS. DNA was isolated from FCG hippocampi (n=3/group). Methylation in CpG (CG) and non-CpG (CH) contexts was assessed by WGoxBS (n=3/group). **A)** There was no main effect difference or interaction in overall whole genome (WG) levels of methylation in CG context (mCG) (Two-way ANOVA). **B)** XX mice had higher levels of X-chromosome mCG than XY mice regardless of gonadal sex (M v. F) (Two-way ANOVA, main effect sex chromosome complement, ***p<0.001). There was also a significant interaction of gonadal and chromosomal sex (p=0.02). **C**) Within repetitive elements of the X-chromosome there was higher mCG in gonadal females than males, with no effect of sex chromosome (Two-way ANOVA, main effect gonadal sex, #p<0.05) and no interaction. **D**) Within non-repetitive elements of the X-chromosome, there was higher mCG in XX genotypes as compared to XY, regardless of gonadal sex (Two-way ANOVA, main effect sex chromosome complement, ***p<0.001) with no significant interaction between gonadal and chromosomal sex. **E)** There was no main effect difference or interaction in genome-wide methylation in non-CpG context (mCH) (Two-way ANOVA). **F)** XX mice had lower mCH than XY mice specifically on the X-chromosome (Two-way ANOVA, main effect sex chromosome complement, ***p<0.001), regardless of their gonadal sex and with no significant interactive effect. The differences in X-chromosome mCH are seen in both (**G**) repetitive and (**F**) non-repetitive elements of the genome. mCG levels were calculated with respect to genic regions by binning 200 nucleotides in flanking regions and region-size dependent bins within the genic region (CGI, gene body, and CTCF) as to maintain the same number of bins for each feature. The average for all (**I**) CGI, (**J**) Gene Body, and (**K**) CTCF were assessed for each of the FCG (XXF, XXM, XYF, XYM) and plotted as averages with 95% CI. **I)** X-chromosome CpG Islands (CGI), shores, and shelves have higher levels of mCG in XX genotypes as compared to XY. The greatest difference in mCG is in the CGI. **J)** X-chromosome gene bodies and flanking regions (+/- 4 Kb) have higher levels of mCG in XX genotypes as compared to XY. The greatest difference in mCG is upstream of TSS (i.e. promoter region). **K)** X-chromosome CTCF binding sites have higher levels of mCG in XX genotypes as compared to XY. The difference in mCG (XX-XY) is greater within the CTCF binding site than in flanking regions (+/- 4 Kb).

When focused on the X chromosome alone, XX genotypes had higher mCG levels than XY genotypes, regardless of gonadal sex (M v. F) (**Fig. 3B**; Two-way ANOVA, main effect sex chromosome complement (XX v. XY), ***p<0.001), with a significant interactive effect of chromosomal and gonadal sex (p=0.0206). The difference in X-chromosome mCG load between XX and XY genotypes was primarily driven by non-repeat regions (**Fig. 3D**; Two-way ANOVA, main effect sex chromosome complement (XX v. XY), ***p<0.001), and was not observed in repetitive elements (**Fig. 3C**; Two-way ANOVA). However, there was a small, but significant effect of gonadal sex on the average mCG in X-chromosomal repetitive elements (Two-way ANOVA, main effect gonadal sex, #p<0.05).

In non-CpG (CH) context, there was no genome-wide difference in methylation (**Fig. 3E**). However, there were higher levels of X-chromosome mCH in XY than XX, regardless of gonadal sex (**Fig.3F;** Two-way ANOVA, main effect sex chromosome complement (XX v. XY), ***p<0.001). Conversely to mCG, higher mCH was seen in both repetitive (**Fig. 3G**) and non-repetitive (**Fig. 3H**) elements of the X-chromosome in XY genotypes as compared to XX (Two-way ANOVA, main effect sex chromosome complement (XX v. XY), ***p<0.001), regardless of gonadal sex, with no interaction of gonadal and chromosomal sex. These X-chromosomal methylations trends are consistent with previous reports from wild-type C57Bl/6 mice [75].

Since the sex-chromosomally driven (XX v. XY) difference in X chromosomal mCG appeared to be concentrated in non-repetitive elements of the X chromosome, we assessed the mCG patterning in and around important gene regulatory regions, including CpG islands, gene bodies, and CTCF binding sites. CpG islands (CGIs) are relatively long stretches (500-2000nt) of GC-rich DNA that are predominantly unmethylated [76]. CGIs have higher methylation on the Xi [77]. In the FCG hippocampi, XX genotypes have higher levels of mCG within CGIs, shores, and shelves than XY genotypes, regardless of gonadal sex. The largest average mCG difference occurred in the CGI followed by the shores and then shelves (**Fig. 3I**; Two-way ANOVA, main effect chromosome (XX v. XY)). While on average most CGIs are hypermethylated in XX (over XY) genotypes, there was a small subset of genes which show hypomethylation of CGIs in XX (vs. XY).

Similarly, gene bodies and the regions 4 kilobases upstream from the transcription start site (TSS) and 4 kb downstream of the transcription end site (TES) had higher percentage of mCG in XX than in XY genotypes (**Fig. 3J**; Two-way ANOVA, main effect chromosome (XX v. XY)). The largest difference in mCG was in the regions 4 kilobases upstream from the transcription start site (TSS), inclusive of the gene promoter.

CCCTC-binding factor (*CTCF*) is a zinc-finger protein that mediates chromatin insulation and gene expression by binding 12- to 20-bp DNA motifs (CTCF binding sites) and altering the 3- dimensional chromatin structure. CTCF has high affinity for certain RNA transcripts, including *Xist* and anti-sense transcript *Tsix* [78] which may help to differentially package the inactive and active X chromatin. In the FCG hippocampus, there was lower percentage of mCG in X Chromosome CTCF binding sites as compared to mCG on the whole X chromosome in all genotypes (XXF, XXM, XYF, XYM). There were higher mCG levels in XX hippocampi than XY, regardless of gonadal sex (**Fig. 3K**). The magnitude of difference between XX and XY mCG in CTCF binding sites (∼10%) was much greater than the average difference seen across the X chromosome (∼3%). Together, the methylation analysis of the X chromosome in FCG hippocampi suggests that X chromosome methylation in non-repetitive regions is regulated by sex chromosome complement (XX v. XY) and likely not influenced by gonadal status.

### X-chromosomal repeat element methylation

Within repetitive regions of the X-chromosome, the regulation of DNA methylation is complex. Here we present evidence that gonadal sex impacts X-chromosomal repetitive element DNA methylation and suggest a hormonally-driven mechanism that may contribute to the gonadal sex effect seen in X-chromosomal repetitive element DNA methylation. The mouse X chromosome is made up of ∼35% LINE-1 (L1) repeats, as opposed to ∼20% on autosomes. The “Lyon Repeat Hypothesis” [79] proposed that LINE-1 (L1) elements promote X-chromosome heterochromatization during XCI. In the FCG hippocampus, LINE-1 elements had higher methylation than LINE-2 (L2) and SINE elements (Alu, B2, B4, MIR) (**Fig. 4A**). Interestingly, the average mCG across X-chromosomal L1 elements was higher in XY genotypes than XX (Two-way ANOVA, main effect sex chromosome complement (XX v. XY), ***p<0.001), indicating that there might be some L1 elements that are not entirely silenced on the inactive X.

**Fig. 4.**
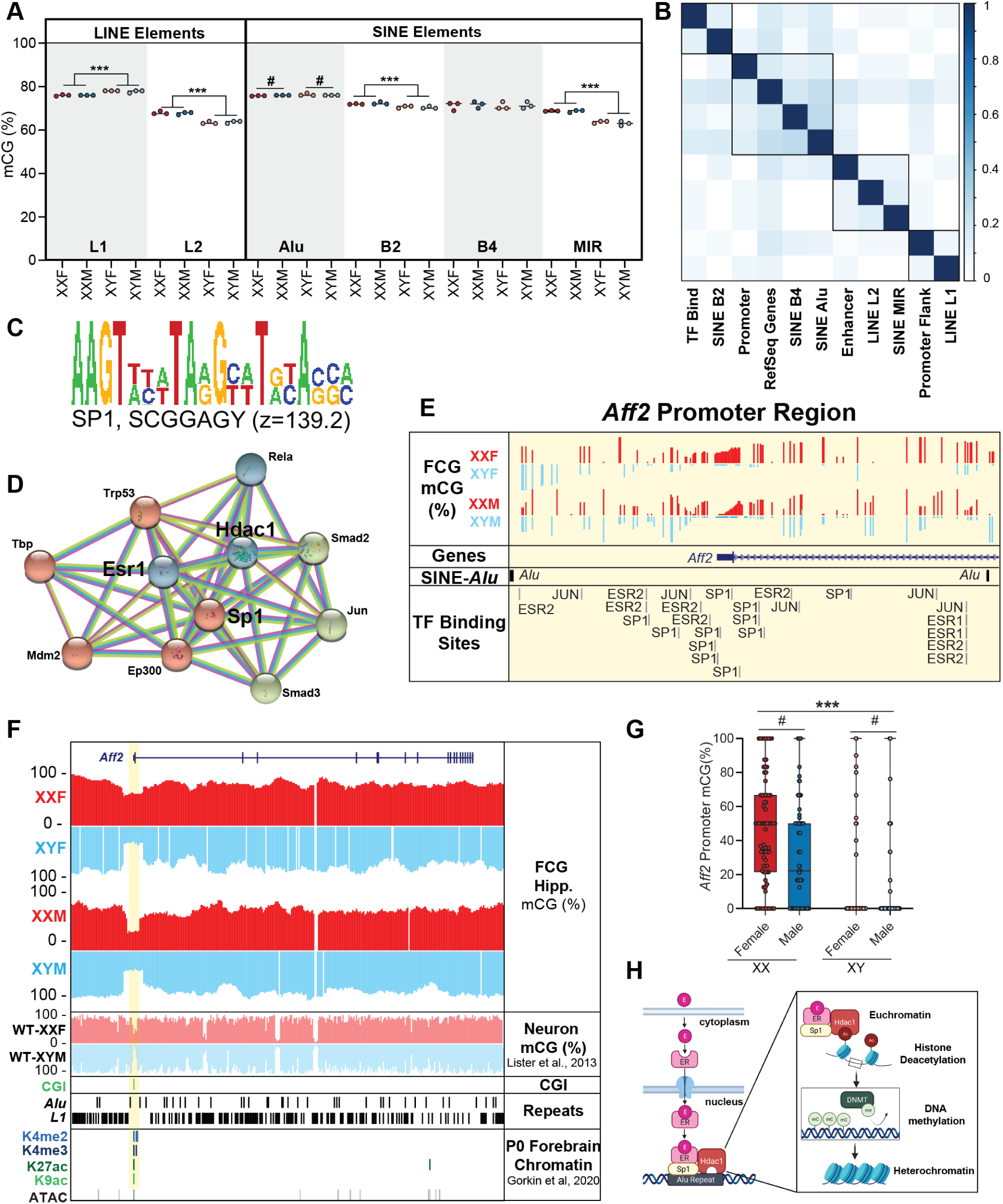
X-chromosome repetitive element DNA methylation in adult FCG hippocampus. FCG DNA methylation levels in X-chromosomal repetitive elements were assessed by WGoxBS (n=3/group). **A)** Average mCG in common X-chromosomal LINE (L1, L2) and SINE (Alu, B2, B4, MIR) repetitive elements were compared between FCG groups for main effects of chromosomal and gonadal sex, as well as interactive effects of gonadal and chromosomal sex. LINE-L1 elements had higher mCG in XY genotypes as compared to XX, regardless of gonadal sex (Two-way ANOVA, main effect sex chromosome complement, ***p<0.001). LINE-L2, SINE-B2, and SINE-MIR repetitive elements had higher mCG levels in XX genotypes as compared to their XY counterparts, regardless of gonadal sex (Two-way ANOVA, main effect sex chromosome complement, ***p<0.001). SINE-Alu repetitive elements had higher methylation in gonadal females than in gonadal males (Two-way ANOVA, main effect gonadal sex, #p<0.05), and a significant interactive effect of gonadal and chromosomal sex (p=0.0231). There was no difference in mCG in SINE-B4 elements. **B)** X-chromosomal browser extensible data (BED) files for common gene regulatory regions and repetitive elements were correlated by Jaccard distance using the BedSect application (Mishra et al., 2020). **C)** TrawlerWeb motif analysis (Dang et al., 2018) on X-chromosomal SINE-*Alu* repeat elements identified SP1 as a potential transcription factor interactor. **D)** STRING protein network analysis (von Mering et al., 2005) of SP1 identified a network of proteins that interact with SP1, including estrogen receptor 1 (ESR1) and histone deacetylase 1 (HDAC1). **E)** *Aff2* (DEG by gonadal sex in Fig. 2F) promoter genome tracks including representative mCG from FCG hippocampus (present study) compared to SINE-*Alu* and transcription factor binding sites (SP1, ESR1, ESR2, JUN) distribution. **F)** *Aff2* gene body genome tracks including representative mCG from FCG hippocampus (present study) compared to genomic features (CGI, SINE-Alu, LINE-L1), previously published DNA methylation from male and female neurons (Lister et al., 2013), and previously published chromatin data from P0 forebrain (Gorkin et al., 2020). **G)** DNA methylation at individual CG sites within the promoter region of *Aff2* bound by the flanking SINE-Alu repeats (highlighted yellow in **Fig. 4E-F**) were compared between FCG groups. *Aff2* promoter methylation was higher in XX genotypes compared to XY (Two-way ANOVA, main effect sex chromosome complement, ***p<0.001) and higher in females than males (main effect gonadal sex, *#p<0.001), as well as an interactive effect of gonadal and chromosomal sex (p=0.001). **H)** Working model for gonadal effect on X-chromosomal methylome. We hypothesize that circulating estradiol binds to ESR1, which then enters the nucleus. The estrogen receptor forms interactions with SP1 and HDAC1, which then binds to X-chromosomal SINE-Alu repeats. The HDAC1 then removes active histone acetylation marks and recruits DNA methyltransferases to hypermethylate the region and compact the chromatin. Created with BioRender.com.

In contrast, X-chromosomal LINE-L2, SINE-B2, and SINE-MIR elements have lower mCG in XY genotypes as compared to their XX counterparts (**Fig. 4A**; Two-way ANOVA, main effect sex chromosome complement (XX v. XY)), consistent with differences seen in whole X-chromosomal methylation. SINE-*Alu* repeats were the only repetitive element examined that showed a main effect of gonadal sex on X-chromosomal mCG levels, with higher methylation in females than in males (**Fig. 4A**; Two-way ANOVA, main effect gonadal sex (M v. F), #p<0.05). SINE-B4 elements showed no difference in mCG by chromosomal or gonadal sex.

Correlation of the degree of overlap (Jaccard distance) between LINE (L1, L2) and SINE (Alu, B2, B4, MIR) repeat elements on the X-chromosome with important gene regulatory features (Promoters, Gene Bodies, Enhancers, Promoter flanks, Transcription factor binding sites) revealed that SINE-Alu and SINE-B4 elements were most strongly correlated with gene promoters and bodies (**Fig. 4B**). L1 elements were associated with promoter flanks and L2 with enhancers, suggesting that LINE elements serve distal regulatory functions in X-inactivation, perhaps through chromatin looping [80]. On the other hand, SINE elements seem tightly correlated with proximate gene regulatory factors, including promoters and transcription factor binding sites. We hypothesize that, in addition to serving as boundary for heterochromatic domains, SINE-*Alu* elements serve as a template to recruit epigenome modifiers to regulate gene transcription (especially in genes that escape X-inactivation) in response to gonadal hormone signals. Motif analysis of X-chromosomal SINE-*Alu* repeat sequences identified SP1 as a potential transcription factor related to these sequences (**Fig. 4C**). STRING protein network analysis [81] identified estrogen receptor 1 (ESR1) and histone deacetylase 1 (HDAC1) as predicted interactors to SP1 (**Fig. 4D**).

In the adult FCG hippocampus, *Aff2* was differentially expressed by gonadal sex (M v. F), with higher expression in gonadal males as compared to females (**Fig. 1F**), regardless of sex chromosome complement. Investigation of the *Aff2* promoter region revealed TSS flanking by SINE-*Alu* repeats, as well as transcription factor binding sites for SP1, ESR1, ESR2, and JUN (**Fig. 4E**). To better understand the epigenomic regulation of *Aff2*, we examined representative genome tracks of FCG hippocampal methylation alongside publicly available methylation and chromatin data, as well as positioning of CpG islands (CGI) and L1/Alu repeats (**Fig. 4F**). *Aff2* has a CGI-containing promoter that coincides with active histone marks (H3K4me2, H3K4me3, H3K27ac, H3K9ac) and open chromatin (ATAC-Seq) peaks from P0 forebrain [82]. Flanking the *Aff2* TSS are *Alu* repeats (both up and down stream), with L1 elements completely absent from the promoter region, but densely populating the region up and downstream of the flanking *Alu* repeats. Considering only CpG sites between the two *Alu* repeats flanking the TSS, we quantified the site-specific methylation within each of the FCG groups. Within this region, mCG was higher in XX compared to XY (**Fig. 4G;** Two-way ANOVA, main effect sex chromosome complement, ***p<0.001) and higher in gonadal females compared to males (**Fig. 4F**; Two-way ANOVA, main effect gonadal sex, #p<0.001), as well as a significant interaction between chromosomal and gonadal sex (p=0.001). Consistent with these results, the *Aff2* promoter was hypomethylated in WT males (compared to female) [83] mirroring the patterning seen in the FCG hippocampus. These data suggest that gonadal sex (and potentially gonadal hormones) may contribute to X-chromosomal DNA methylation and have implications in escape from X-inactivation. Based on the associations observed here, we propose that circulating estradiol binds to ESR1 and in the nucleus complexes with SP1 and HDAC1. The complex interacts with the SINE-*Alu* repeats flanking active promoters to induce changes in the chromatin landscape, including hypermethylation and heterochromatization (**Fig. 4H).**

### Epigenomic patterning in genes that consistently escape X-chromosome inactivation in mouse hippocampus

There were four X-chromosomal genes (*Kdm6a*, *Ddx3x*, *Eif2s3x*, *Xist*) that were found to consistently escape X-inactivation across all previously published hippocampal transcriptomic studies examined in **Fig. 2A** and in the present FCG hippocampal study. *Kdm6a*, *Ddx3x*, and *Eif2s3x* are expressed in all FCG groups (XXF, XXM, XYF, XYM) and wildtype (WT-XXF, WT-XYM) with higher expression in XX genotypes regardless of gonadal sex, whereas *Xist* expression is only detected in FCG XXF, FCG XXM, and WT-XXF. Thus, we consider the epigenomic regulation of *Kdm6a*, *Ddx3x*, and *Eif2s3x* separate from *Xist*.

Examining the genome tracks for *Kdm6a*, *Ddx3x*, and *Eif2s3x*, revealed several epigenomic patterns across these genes that consistently escape X-inactivation in the mouse hippocampus (**Fig. 5A**). Namely, all three escapees have promoters that contain large CGIs (>500 nt) and coincide with active histone marks (H3K4me2, H3K4me3, H3K27ac, H3K9ac) from publicly available CNS data [82]. Further, the promoter region is almost entirely unmethylated in FCG hippocampus (regardless of chromosomal or gonadal sex) and WT neurons (both males and females) [83] (**Fig. 5A**). The escapee gene bodies each contained H3K36me3 marks in previously published CNS data [82], although H3K36me3 peaks only cover 0.25% of the X-chromosome. Each or the three escapee promoters are flanked by SINE-*Alu* repeats (+/- 5kb from TSS) and are relatively depleted in intragenic LINE-L1 elements (<6% L1) compared to the whole X-chromosome (∼33% L1).

**Fig. 5.**
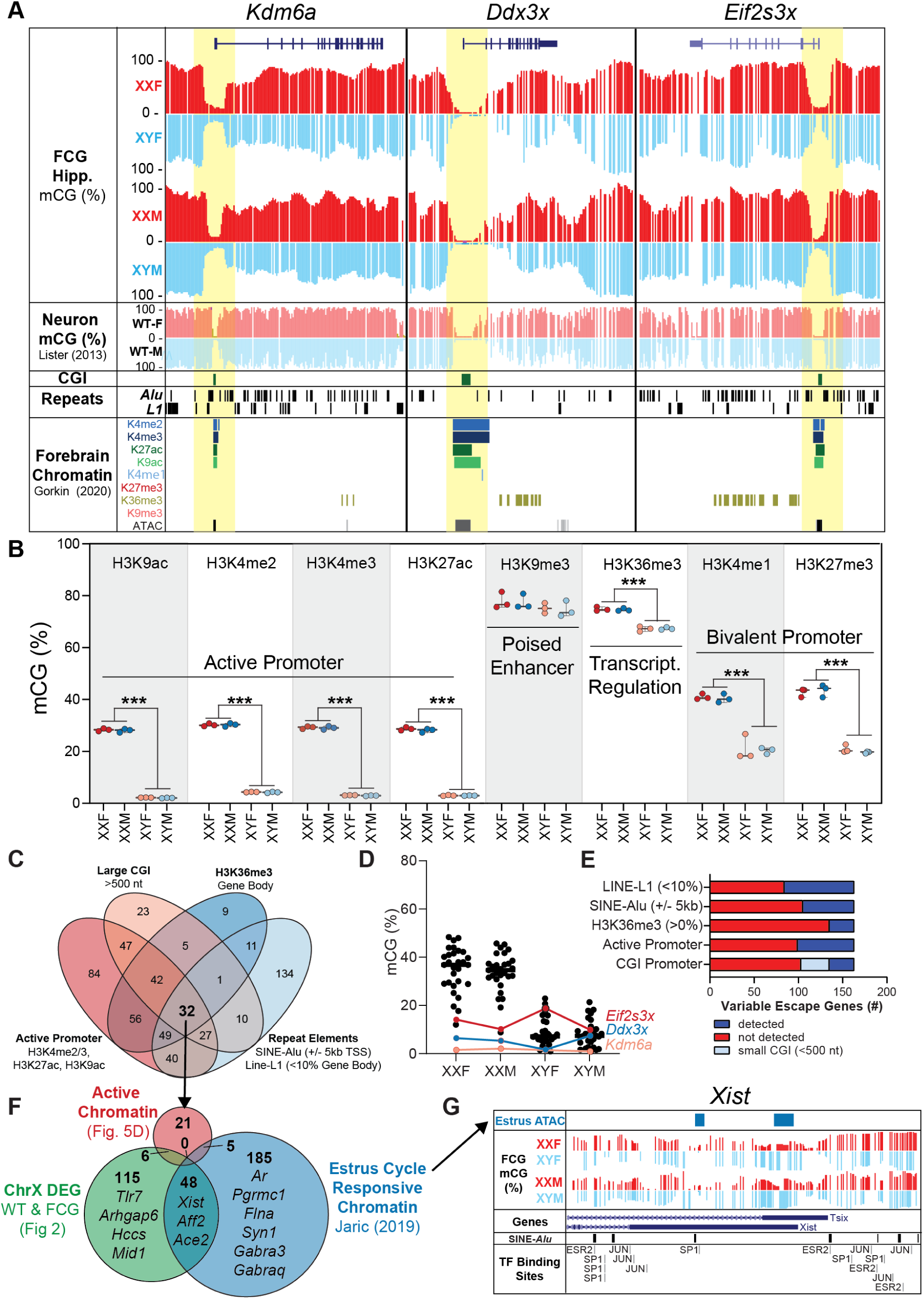
Epigenomic signatures of genes that escape X-inactivation. FCG hippocampal DNA methylation levels within different genomic contexts were compared for genes that escape X-inactivation. **A)** *Kdm6a*, *Ddx3x*, and *Eif2s3x* were found to consistently escape X-inactivation in mouse hippocampus from the previous studies examined in Fig. 2a and FCG (present study). Genome tracks for *Kdm6a*, *Ddx3x*, and *Eif2s3x* compared FCG mCG patterning to genomic features (CGI, Repeats) and previously published CNS mCG (Lister et al., 2013) and chromatin (Gorkin et al., 2020) patterns. **B)** Average methylation across published CNS X-chromosomal histone modification peaks (Gorkin et al., 2020), showed higher methylation in XX genotypes as compared to XY genotypes in H3K9ac, H3K4me2, H3K4me3, H3K27ac, H3K36me3, H3K4me1, and H3K27me3 regardless of gonadal sex, with no interactive effect (Two-way ANOVA, main effect sex chromosome complement, ***p<0.001). There was no detected difference in mCG in H3K9me3 peaks. **C)** Commonalities between the chromatin structure in genes that consistently escape X-inactivation include: 1) Large CGIs (>500 nt) within the promoter, 2) Active promoter histone marks (H3K4me2, H3K4me3, H3K27ac, H3K9ac), 3) H3K36me3 gene body peaks, 4) SINE-*Alu* flanks (+/- 5kb from TSS), and 5) low LINE-L1 gene body (<10%). There were 32 genes which had the same chromatin signature as the common escape genes (**Supplemental Table X**). **D)** Promoter mCG (%) for the 32 genes with the same chromatin signature as the escape genes. All genes, except *Eif2s3x, Ddx3x,* and *Kdm6a* (common escape), had higher mCG in XX genotypes compared to XY genotypes (Two-way ANOVA, main effect sex chromosome complement, ***p<0.001), regardless of gonadal sex with no interaction. **E)** There were 169 X-chromosomal genes that show variable escape from X-inactivation (i.e. differentially expressed in mouse hippocampus by gonadal and/or chromosomal sex in the previous studies examined in Fig. 2A and/or the present FCG study). These 169 variable escape genes were examined for their concordance with the five signatures of consistent escape gene chromatin. The majority of variable escape genes did not have chromatin signatures consistent with the consistent escape genes. **F)** Comparison of 32 genes that share the active chromatin signature from Fig. 5C, the 169 sex differentially expressed genes from Fig. 5E, and 238 genes with estrus cycle-responsive chromatin (Jaric et al., 2019), identified 48 differentially expressed genes that have estrus-responsive chromatin. **G)** *Xist/Tsix* locus genome tracks showing overlap with estrus-responsive chromatin (Jaric et al., 2019), SINE-*Alu* repeats and SP1, ESR2, and JUN transcription factor binding sites.

Within the X-chromosomal transcriptomic analysis, we identified histone H3-K4 and H3-K27 demethylation, as over-represented pathways in X-chromosomal sex differences. X-encoded histone modifiers *Kdm6a* and *Kdm5c* demethylate H3K27me2/3 [84] and H3K4me2/3 [85], respectively. The permutation of histone modifications delineates important gene regulatory regions, including active promoters (H3K27ac, H3K9ac, H3K4me2, H3K4me3), weak/inactive promoters (H3K9ac, H3K4me3, H3K4me2, H3K4me1), bivalent promoters (H3K27me3, H3K4me3, H3K4me2, H3K4me1), and transcriptional regulation (H3K36me3). The publicly available CNS histone modification data used for comparison was from pooled male and female brains at postnatal day 0 [82] and serves as a mixed sex developmental baseline. We then use differences in DNA methylation within the histone peaks to predict sex differences in the histone modification patterns (which may be influenced by female-biased *Kdm6a* and *Kdm5c* activity).

Within marks of active promoters (H3K9ac, H3K4me2, H3K4me3, H3K27ac) there was higher mCG in XX as opposed to XY genotypes, regardless of gonadal sex (**Fig. 5B**). Since promoter hypermethylation is generally associated with gene inactivation [86], this difference is likely a primary mechanism in maintenance of X-inactivation. As such, hypermethylation of XX active promoters, is likely accompanied by a transition to an inactive promoter state which could involve H3K4 demethylation by *Kdm5c*. Hypermethylation of gene body DNA coincides with H3K36me3 and is proposed to prevent cryptic transcription from non-canonical transcription start sites (TSSs) [87]. Hypermethylation of H3K36me3 loci in XX genotypes compared to XY (**Fig. 5B**, Two-way ANOVA, main effect sex chromosome complement, ***p<0.001) could indicate demarcation of X-inactivation escape, as previously reported [88]. Repressive H3K4me1 and H3K27me3 are often found at bivalent promoters along with active H3K4me3 [89]. Both H3K4me1 and H3K27me3 peaks showed higher methylation in XX over XY genotypes (**Fig. 5B**, Two-way ANOVA, main effect sex chromosome complement, ***p<0.001). Since H3K4me1 and H3K27me3 are generally repressive marks, we interpret the XX hypermethylation as indication that XX genotypes have higher H3K4me1 and H3K27me3 within their promoter regions, a function of X-inactivation. As an H3K27 demethylase, *Kdm6a* is likely responsible for liberating repressed chromatin to allow escape from X-inactivation at specific genes.

To summarize, differences in DNA methylation within X-chromosomal histone modifications established early in the brain [82] are regulated entirely by sex chromosome complement (XX v. XY), with no effect of gonadal sex. The differences in mCG are most likely associated with variance in histone modification between XX and XY, induced (in part) by imbalance in X-encoded histone modifying enzymes.

In addition to genes that always escape X-inactivation in the mouse hippocampus (i.e. *Kdm6a*, *Ddx3x*, *Eif2s3x*), there are also several genes that variably escape X-inactivation; potentially depending on age, disease state, and interaction with gonadal hormones. To determine if genes with similar epigenomic landscapes had similar escape expression patterns, we compiled a list of 32 X-chromosomal genes meeting the following criteria: 1) large CGI (>500 nt), 2) active promoter (H3K27me2, H3K27me3, H3K9ac, H3K27ac), 3) H3K36me3 gene body methylation, 4) SINE-*Alu* (+/- 5kb TSS), and 5) *L1* depleted in gene body (<10%) (**Fig**. **5C; Supplemental Table 4**). However, unlike the escape genes which showed almost entirely unmethylated promoter regions across all groups (with no difference by chromosomal or gonadal sex), the other 27 of the 29 identified genes had higher methylation in XX genotypes, as compared to XY genotypes (**Fig. 5D,** Two-way ANOVA, Bonferonni MTC, p<α=0.0016, main effect sex chromosome complement). Two genes (*Rlim*, *Fam50a*) with the escape chromatin signature had no difference in promoter mCG. However, the promoters mCG of *Rlim* (∼28%) and *Fam50a* (∼29%) were higher than the consistent escape genes (∼6%). Thus, it seems that protecting the Xi promoter from methylation is necessary for consistent escape. In addition, the methylation levels of XXF/M (∼32%) and XYF/M (∼8%) are indicative of potential shifting between active and poised chromatin.

To determine how the chromatin structure of 160 variably escaping genes (identified as X-chromosomal DEG in previous WT [54–57] or present FCG studies) (**Supplemental Table 2-3**), we compared the number of genes that met each of the five escape chromatin criteria (**Supplemental Table 4)**. We found that only three variable escape genes (*Med14, Magt1, BC065397*) shared the same chromatin signature as the common escape genes (*Kdm6a, Eif2s3x,* and *Ddx3x*). However, the promoter regions of these genes (**Supplemental Table 4**), are hypermethylated in XX genotypes compared to XY, suggesting that the epigenomic landscape (including histone marks) are likely different between the active and inactive X.

Of the variable escape genes: 27% had large CGI promoters, 39% had active promoter histone marks, 17% had gene body H3K36me3, 55% had SINE-*Alu* promoter flanks, and 49% had low L1 gene body density (<10%) (**Fig. 5E**). Thus, it seems likely that the epigenomic regulation of variably escaping genes is distinct from that of common escape genes. As previously suggested, we believe that X-chromosomal DNA methylation could be modulated by gonadal hormone levels. ATAC-Seq profiling of the female brain across the estrus cycle identified 238 genes closest to estrus-responsive chromatin [90]. Intersecting the 238 estrus-responsive chromatin genes with the 160 variable escapee genes and the consistent escape chromatin signature, identified 48 estrus-responsive genes with variable escape and a chromatin signature distinct from the consistent escape genes (**Fig. 5F, Supplemental Table 4**). *Xist* was one of the estrus-responsive DEGs and examination of the *Xist*/*Tsix* locus revealed estrus-responsive chromatin peaks in the promoter of *Xist* and transcription end site of *Tsix*. Additionally, distribution of SINE-*Alu* repeats mirrored that of TFBS (SP1, ESR2, JUN) [91] (**Fig. 5G**). Loss of *Xist* expression (and X-inactivation) in female breast, ovarian, and cervical cancer cell lines [92] further support the premise that hormonal mechanisms may contribute to *Xist* expression.

### Unbiased calling of X-chromosome differentially methylated regions (DMRs) from Whole Genome Oxidative Bisulfite Sequencing (WGoxBS) in FCG hippocampi

To assess FCG X-chromosomal methylation patterning in an unbiased fashion, we called X-chromosomal mCG DMRs with 1 kb non-overlapping windows and minimum average difference of 10% between at least two groups (Chisq-test, sliding linear model (SLIM) q<0.05), followed by a Two-way ANOVA to determine main effects of chromosomal and gonadal sex, as well as potential interactions (Bonferroni correction for six pairwise comparisons, p<α=0.0083). Using these criteria, we identified 2552 DMRs: 2456 by sex chromosome (XX v. XY), 145 by gonadal sex (M v. F), and 87 interactive effects (**Fig. 6A, Supplemental Table 5**) on the X-chromosome. Consistent with our previous findings (**Fig. 3-5**), sex-chromosomally regulated DMRs were: 1) enriched in gene regulatory features (gene body, promoter, TFBS, enhancer, CGI, CTCF) and CNS histone marks for active promoters (H3K4me2, H3K4me3, H3K27ac, H3K9ac), poised promoters (H3K4me1, H3K27me3) and transcriptional regulation (H3K36me3), and 2) depleted in repetitive elements (LINEs, SINEs, LTRs) (Fisher’s exact test, p<0.05) (**Fig. 6B**). Correlation of Jaccard distances of the assessed genomic features with sex chromosomally-driven DMRs, show strong correlation to active promoter regions, again suggesting strong sex chromosome autonomous regulation of X-inactivation (**Fig. 6C**). Within unexpressed genes in the FCG hippocampi there was a small but significant difference in promoter methylation between XX (∼68%) and XY (∼64%) (**Fig. 6D**, Two-way ANOVA, main effect sex chromosome complement, ***p<0.001), with no effect of gonadal sex. Within expressed gene promoters, there was a large difference between XX (∼40%) and XY (∼17%) genotypes (**Fig. 6D**, Two-way ANOVA, main effect sex chromosome complement, ***p<0.001), with no effect of gonadal sex. The incidence of CGI-containing promoters was much higher within expressed genes (∼61%) as compared to unexpressed genes (∼11%) (Chi-sq test, p<0.05) (**Fig. 6E**). There was a strong, negative association between the difference in promoter mCG (XX-XY) and log(FC(XX/XY)) gene expression (Pearson r, p<0.05) with no difference in slope or intercept of the linear fit between gonadal males and females. X-chromosomal genes with XX-biased expression (XX>XY) had smaller differences in promoter methylation (XX-XY), while genes with XY-biased genes had larger differences in promoter methylation (XX-XY). These results are consistent with the escape signature of *Kdm6a, Ddx3x*, and *Eif2s3x* (**Fig. 5A**).

**Fig. 6.**
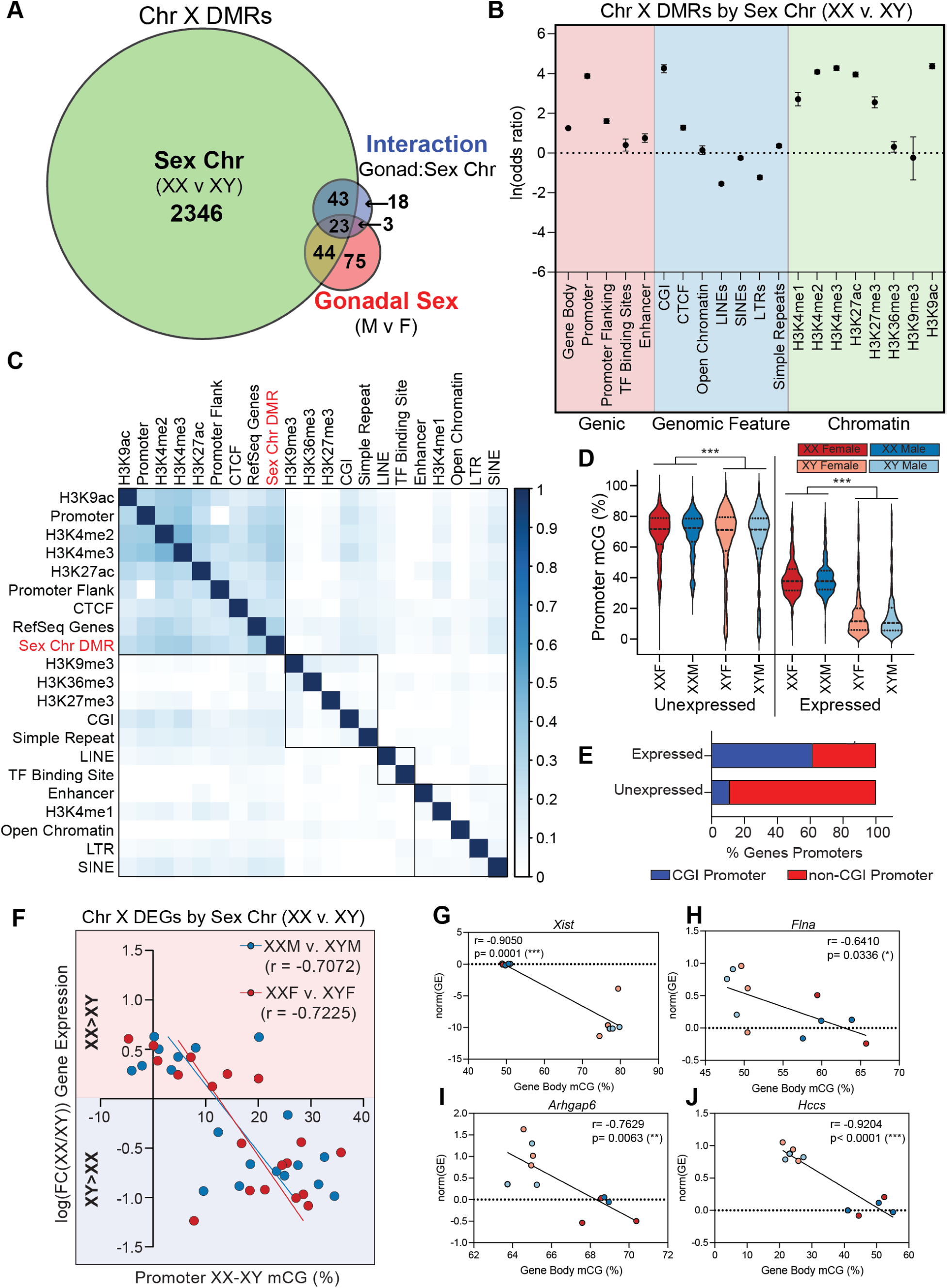
Epigenomic analysis of X-chromosomal differentially methylated regions (DMRs) from adult FCG hippocampi. DNA was isolated from FCG hippocampi (n=3/group). Methylation in CG context was assessed by WGoxBS (n=3/group) and analyzed in methylKit to identify ChrX DMRs (w=1000). ChrX DMRs with minimum mCG difference of 10% between at least two groups (Chisq-test, sliding linear model (SLIM) q<0.05) were assessed for main effects of sex chromosome (XX v. XY) and gonadal sex (M v. F) (2-way ANOVA, Bonferonni α<0.0083). **A)** Venn diagram of DMRs by sex chromosome (XX v. XY), gonadal sex (M v. F), and interactive effect of gonadal and chromosomal sex. **B)** Relative enrichment and depletion of ChrX DMRs by sex chromosome (XX v. XY) within various genomic features. **C)** BedSect (Mishra et al., 2020) correlation plot of ChrX genomic regulatory features (Ward hierarchical clustering, Jaccard similarity index). **D)** Distribution of unexpressed and expressed gene promoter mCG (%). Unexpressed gene promoters (+/- 2kb from TSS) had higher mCG in XX genotypes (68%) as compared to XY genotypes (54%), regardless of gonadal sex. Expressed gene promoters had higher mCG in XX genotypes (40%) as compared to XY genotypes (17%), regardless of gonadal sex (2-way ANOVA, main effect sex chromosome complement, ***p<0.001). **E)** Only 11% of unexpressed ChrX gene promoters contain CGIs, as compared to 61% of expressed ChrX gene promoters. **F)** Correlation of log(FC) gene expression (XX/XY) and difference in gene promoter mCG (XX-XY) for 19 ChrX DEGs identified in Fig. 2E (Pearson r, p<0.05). *Xist* was excluded from the correlation since lack of expression in XY genotypes inflates the fold-change values used for analysis. **G-J)** Intersection of DMRs within gene bodies with differentially expressed genes by sex chromosome (XX v. XY) identified four genes: (F) *Xist,* (G) *Flna*, (H) *Arhgap6*, and (I) *Hccs*, all of which have a significant negative correlation between gene body methylation and gene expression (Pearson r, *p<0.05, **p<0.01, ***p<0.001).

In agreement with the predicate that gene expression anti-correlates with gene promoter methylation, expression of DEGs *Xist* (**Fig. 6G**), *Flna* (**Fig. 6H**), *Arhgap6* (**Fig. 6I**), and *Hccs* (**Fig. 6J**) anticorrelated with gene body methylation of those genes.

Together, these epigenomic analyses suggest that X-chromosomal regulation of gene expression is primarily controlled by sex chromosomal-autonomous mechanisms, reflecting patterns consistent with X-inactivation and escape.

### Targeted bisulfite amplicon sequencing (BSAS) of X-chromosome gene promoters in FCG hippocampal DNA

Our WGoxBS data had 2-6X genome-wide coverage, which is sufficient to analyze methylation values in windows and collapse certain genomic regions. In order to assess base-specific methylation patterning, we performed targeted bisulfite amplicon sequencing (BSAS) within the promoter region of X-chromosome genes, using custom primers (**Table 1**).

**Table 1.**
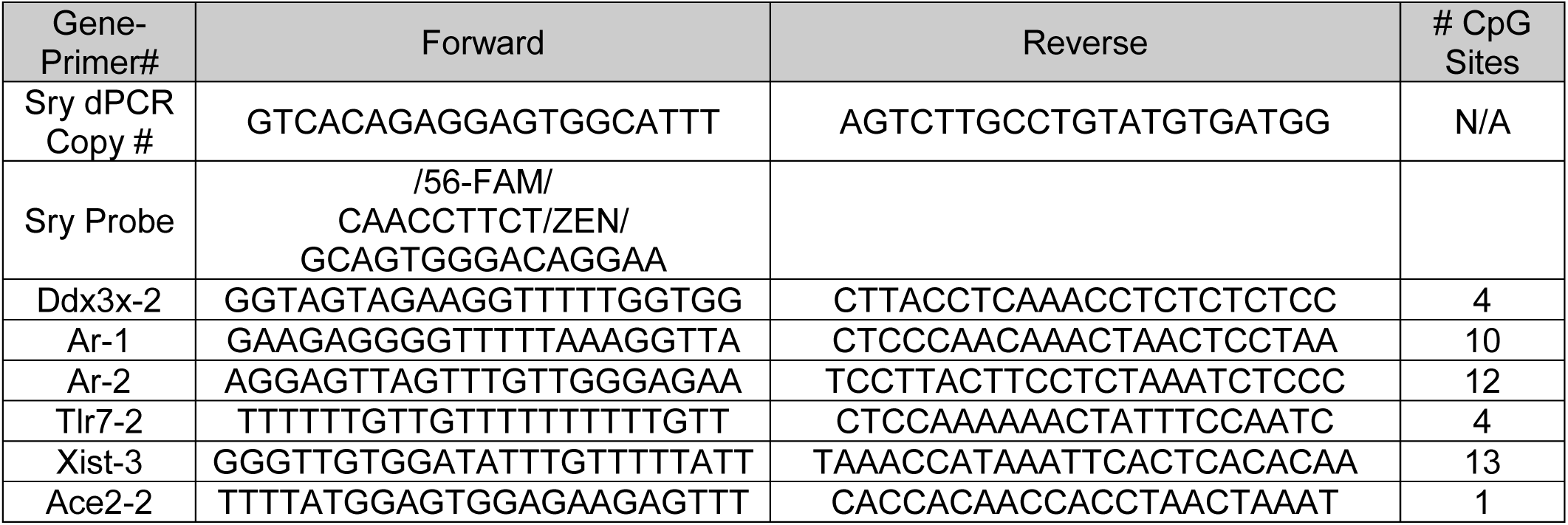
Digital PCR copy number and BSAS primers (5’->3’) for targeted methylation analysis.

X-linked DEAD-box RNA helicase DDX3 (*Ddx3x*) plays an integral role in transcription and translation, as well as splicing and RNA transport with loss-of-function mutations associated with intellectual disability and developmental delays [93]. *Ddx3x* was more highly expressed in XX genotypes (XXF/XXM) as compared to XY genotypes (XYF/XYM). We amplified a region of the *Ddx3x* promoter, containing 4 CpG sites. Average mCG across the region was higher in XX genotypes as compared to XY genotypes, regardless of gonadal sex (**Fig. 7A**), as well as each individual CpG site (**Fig. 7B**). The site-specific methylation appeared to be strongly regulated by sex chromosome complement, as evidenced by the consistent topography of mCG.

**Fig. 7.**
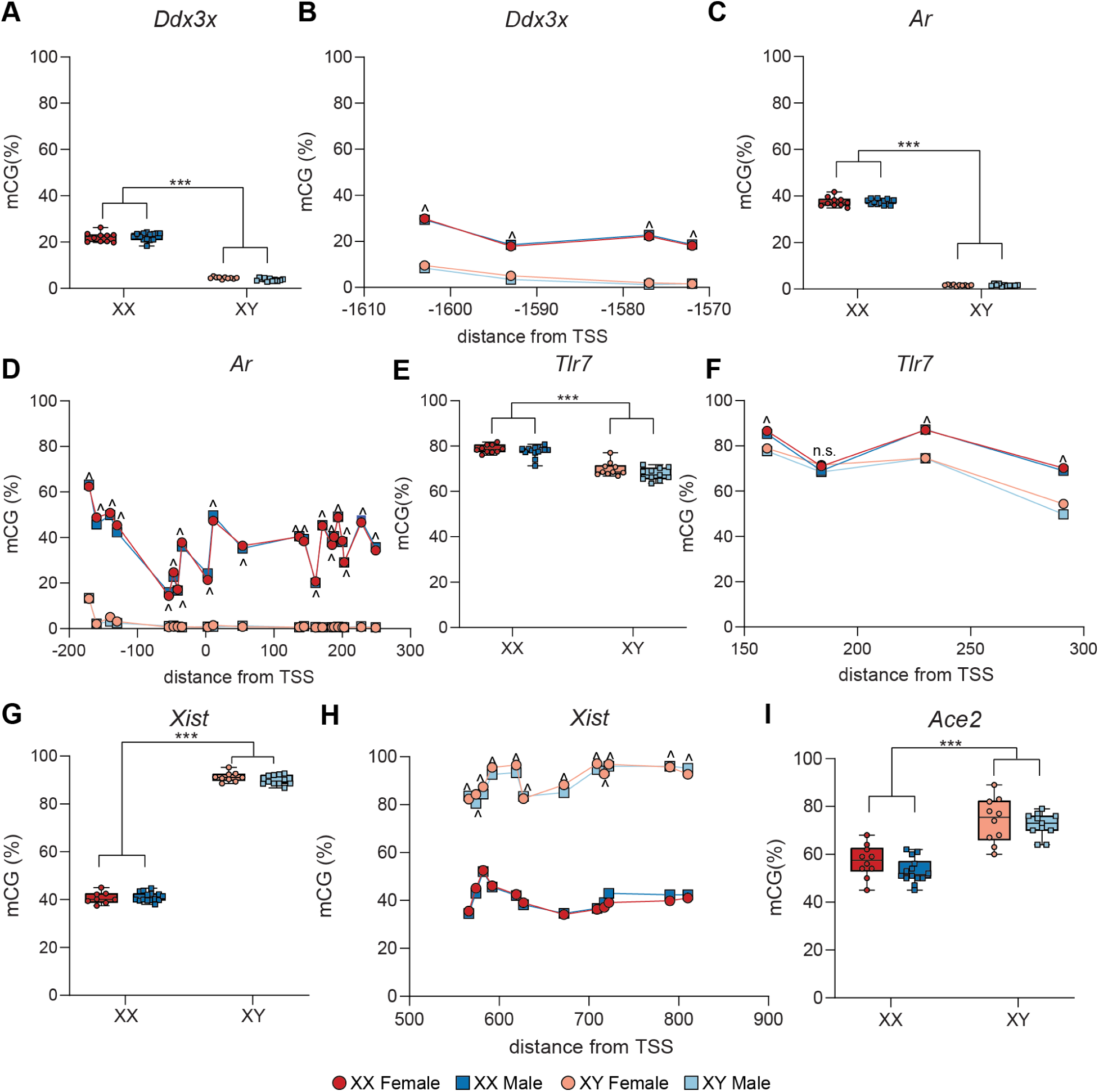
Targeted bisulfite amplicon sequencing (BSAS) of X-chromosome gene promoters in FCG hippocampus. DNA isolated from FCG hippocampi (n=10-16/group) was bisulfite-converted and DNA methylation in gene promoters (+/- 2kb from TSS) was assessed by BSAS. (A) Average *Ddx3x* promoter mCG is lower in XY genotypes (XYF/XYM) as compared to XX genotypes (XXF/XXM), regardless of gonadal sex. (B) The topography, or patterning, of mCG across the amplified region of the *Ddx3x* promoter shows base-specific regulation of mCG by sex chromosome complement (XX v. XY), with higher mCG in XX genotypes than XY genotypes at each CG site in the region. (C) Average *Ar* promoter mCG is lower in XY genotypes (XYF/XYM) as compared to XX genotypes (XXF/XXM), regardless of gonadal sex. (D) The topography of mCG across the amplified region of the *Ar* promoter shows base-specific regulation of mCG by sex chromosome complement (XX v. XY), with higher mCG in XX genotypes than XY genotypes at each CG site in the region. (E) Average *Tlr7* promoter mCG is lower in XY genotypes (XYF/XYM) as compared to XX genotypes (XXF/XXM), regardless of gonadal sex. (F) The topography of mCG across the amplified region of the *Tlr7* promoter shows base-specific regulation of mCG by sex chromosome complement (XX v. XY), with higher mCG in XX genotypes than XY genotypes at three of the four CG sites in the region. (G) Average *Xist* promoter mCG is higher in XY genotypes (XYF/XYM) as compared to XX genotypes (XXF/XXM), regardless of gonadal sex. (H) The topography of mCG across the amplified region of the *Xist* promoter shows base-specific regulation of mCG by sex chromosome complement (XX v. XY), with higher mCG in XY genotypes than XX genotypes at each CG site in the region. (I) Average *Ace2* promoter mCG is higher in XY genotypes (XYF/XYM) as compared to XX genotypes (XXF/XXM), regardless of gonadal sex. **Boxplots** represent median, interquartile range, and minimum/maximum mCG (%) averaged over all CG sites within the amplified region of the respective gene promoter. (2way ANOVA, main effect of sex chromosome complement (XX v. XY), ***p<0.001) (2way ANOVA, Tukey’s *post-hoc*, ^p<0.05 for all four XX v XY comparisons: XXF v XYF, XXF v. XYM, XXM v. XYM, XXM v. XYF)

Following sex-chromosomally driven sex determination, development of the gonads and production of gonadal hormones further drives dichotomization of sexual phenotypes. Testosterone [94] is recognized by the androgen receptor (*Ar*), a hormone nuclear receptor and transcription factor that has many biological functions, including male reproductive organ and secondary sex characteristic development [95]. Androgens have been found to effect hippocampal structure and function, as well as playing a role in hippocampal-dependent behavior, long-term potentiation, and dendritic arborization [96]. *Ar* is an X-chromosomally encoded gene and subject to X-chromosome inactivation. Although *Ar* was not differentially expressed by sex in outside studies (**Supplemental table 2 (Union)**) or in the present study (neither by gonadal sex (M v. F) or sex chromosome complement (XX v. XY)), we wanted to determine if gonadal sex had any effect on *Ar* promoter methylation, as *Ar* was identified within estrus-responsive chromatin (**Fig. 5F**). BSAS analysis of 22 CpGs in the *Ar* promoter region, showed very low (∼0%) mCG in XY genotypes with close to 40% average methylation in XX genotypes, regardless of gonadal sex (**Fig. 7C**). Each CG-site within the amplified region of the *Ar* promoter had lower mCG (∼0%) in XY genotypes compared to XX genotypes (∼10-60%), regardless of gonadal sex. The patterning of *Ar* promoter mCG was well-conserved between XXF and XXM, suggesting tight regulation of *Ar* promoter methylation by sex chromosome complement, with no effect of gonadal sex (**Fig. 7D**).

Toll-like receptor (*Tlr7*) is an X-encoded pattern recognition receptor (PRR), critical in innate immunity. *Tlr7* is primarily expressed on microglia in the brain [97] and initiates a Type I interferon (IFN) response to single-stranded viral RNA (ssRNA) [98]. *Tlr7* was differentially expressed by sex in prior studies (**Supplemental table 2 (Union)**) and by sex chromosome complement in the present study (**Fig. 2E**) with higher expression in XY (v. XX) genotypes. *Tlr7* was not found within estrus-responsive chromatin (**Fig. 5F**). The average mCG in an amplified region of the *Tlr7* promoter containing 4 CG sites is higher in XX genotypes over XY genotypes (**Fig. 7E**), and in three of the four CG sites within the amplified region, with no differences by gonadal sex (**Fig. 7F**).

*Xist* was one X-encoded gene that was differentially expressed by sex in all outside studies examined (**Supplemental table 2 (Intersect)**) and by sex chromosome complement in the present study (**Fig. 2B, 2E**). As a critical regulator of X-inactivation and X-chromosome dosage compensation, we analyzed mCG in an amplified region of the *Xist* promoter in FCG hippocampi. The average mCG within the amplified region of the *Xist* promoter was higher in XY genotypes than XX genotypes, regardless of their gonadal sex (**Fig. 7G**), despite the fact that *Xist* was identified as a gene within estrus-responsive chromatin. The base-specific topography of CG methylation is well conserved by sex chromosome complement (XX/XY), with no effect of gonadal sex (**Fig. 7H**).

Angiotensin-converting enzyme 2 (*Ace2*) is surface receptor responsible for negative regulation of the renin-angiotensin system to modulate blood pressure and fluid/electrolyte balance. *Ace2* recently gained attention as the entry receptor for the novel SARS-coronavirus 2 (SARS-CoV-2) [99]. *Ace2* was differentially expressed by sex in the prior studies we examined (**Supplemental table 2 (Union)**) and by gonadal sex (M v. F) in the present study (**Fig. 2G**), as well as being within close proximity to estrus-responsive chromatin (**Fig. 5F**). We assessed the mCG at a single CpG site within the *Ace2* promoter and found higher mCG in XY genotypes as compared to XX genotypes, irrespective of their gonadal sex (**Fig. 7I**).

In summary, targeted methylation analysis confirmed that X-chromosome methylation within the analyzed regions is tightly regulated, in a base-specific fashion, by sex chromosome complement, and not gonadal sex.

### Chromatin context of ChrX DMRs regulated by gonadal sex

There were 145 ChrX DMRs by gonadal sex, including 67 DMRs that were also differentially methylated by chromosomal sex, and 26 DMRs with significant interactive effects (**Fig. 6A, Supplemental Table 5**). Gonadal sex regulated DMRs were: 1) enriched in gene regulatory features (gene body, promoter, promoter flanking region, CGI, CTCF) and CNS histone marks for active promoters (H3K4me2, H3K4me3, H3K27ac, H3K9ac), poised promoters (H3K4me1) and heterochromatin (H3K9me3) and 2) depleted in LINE repetitive elements (Fisher’s exact test, p<0.05) (**Fig. 8A**). Correlation of Jaccard distances of the assessed genomic features with gonadal sex-driven DMRs, show strong correlation to active promoter regions, CGIs, and estrus-responsive chromatin (**Fig. 8B**).

**Fig. 8.**
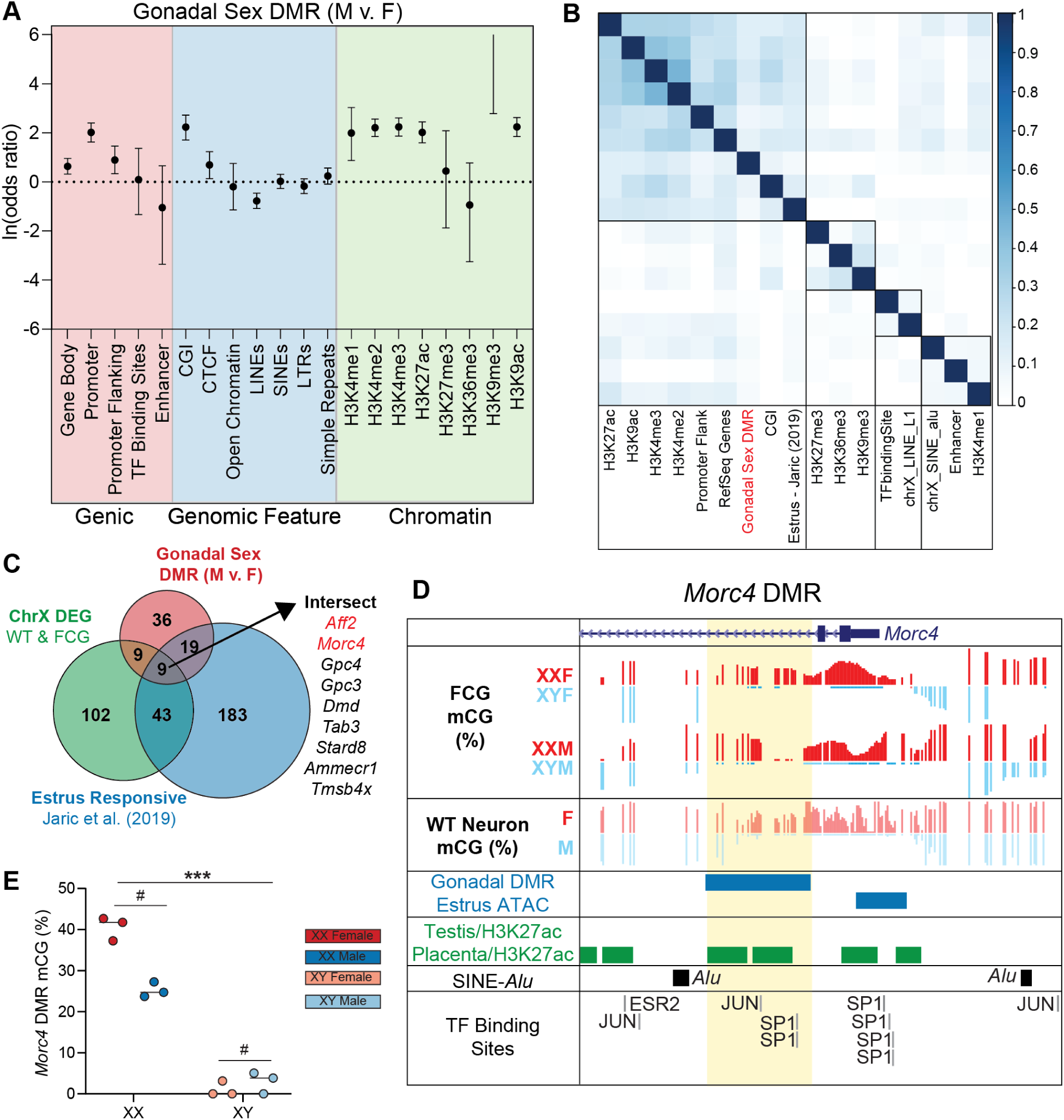
Comparison of DMRs X-chromosome differentially regulated by gonadal sex (M v. F) to chromatin context and gene expression. **A)** Relative enrichment and depletion of ChrX DMRs by gonadal sex (M v. F) within various genomic features. **B)** BedSect (Mishra et al., 2020) correlation plot of ChrX genomic regulatory features (Ward hierarchical clustering, Jaccard similarity index) with gonadal sex DMRs and estrus-responsive chromatin (Jaric et al., 2019). **C)** Comparison of 145 gonadal sex DMRs, 169 sex differentially expressed genes from Fig. 5E, and 238 genes with estrus cycle-responsive chromatin (Jaric et al., 2019), identified 9 DMRs with variable escape from X-inactivation and estrus-responsive chromatin. **D)** Genome tracks of *Morc4* promoter region with gonadal DMR region (highlighted yellow), FCG mCG, WT male and female neuronal mCG (Lister et al., 2013), estrus-cycle differential ATAC-seq peaks (Jaric et al., 2019), male (testis) and female (placenta) specific H3K27ac peaks, SINE-*Alu* repeats, and JASPAR 2022 TFBSs (SP1, JUN, ESR2) (Castro-Mondragon et al., 2022). **E)** *Morc4* DMR mCG (%) is higher in XX vs XY genotypes and higher in M v F, with an interaction between chromosomal and gonadal sex (Two-way ANOVA, main effect sex chromosome complement (***p=1.44E-08), main effect gonadal sex (#p= 0.0011), interaction (p= 0.0002)).

There were 73 X-chromosomal genes that had a gonadal sex DMR within 4kb (+/-) of the gene body. Intersecting these genes with the 160 variable escape genes (**Supplemental Table 2 &3**), and 238 estrus-responsive genes (**Supplemental Table 4**) identified 9 genes (*Aff2, Morc4, Gpc4, Gpc3, Dmd, Tab3, Stard8, Ammerc1, Tmsb4x*) whose escape from X-inactivation may be regulated by sex hormonal mechanisms (**Fig. 8C**). Genome tracks of the *Morc4* promoter region showed that the gonadal DMR (highlighted in yellow) was closely associated with the estrus-responsive ATAC-seq peak [90] and JASPAR TFBS (SP1, JUN, ESR2) [91], was flanked by SINE-*Alu* repeats, and corresponded to sex differential H3K27ac ChIP-Seq peaks (placenta v. testis) [100] (**Fig. 8D**). Together these data suggest that *Morc4* variably escapes from X-inactivation as a result of alterations in gonadal hormones. *Morc4* mCG within the DMR had higher methylation in XX (v. XY) genotypes and higher methylation in gonadal females (v. males) with a significant interaction effect (Two-way ANOVA, main effect sex chromosome complement (***p=1.44E-08), main effect gonadal sex (#p= 0.0011), interaction (p= 0.0002)) (**Fig. 8E**).

### Transcriptomic analysis of autosomal differential expression from adult FCG hippocampi

After establishing that the sex chromosome transcriptome and methylome of FCG hippocampi are primarily controlled by sex chromosome complement (XX v. XY) with some specific interactions with gonadal sex (M v. F), we next examined regulation of autosomal sex differences. We first intersected the previous hippocampal transcriptomic studies to determine steady-state sex differences in the mouse hippocampus (**Fig. 9A**). Although there were no sex differences in common between all studies (**Fig. 9A**, **Supplemental Table 6 (Intersection)**), there were 2896 sex differences identified in at least one study (**Supplemental Table 6 (Union)**). Age, sample size, and estrus-cycle staging differences between these studies are summarized in **Table 2.** To interrogate the discordance in sex differences across these studies, we performed pairwise correlations (Pearson r) on fold change (M v. F) from DEGs expressed in across all studies. Hierarchical clustering of the correlation matrix showed the strongest association between 3 mo [101] and 4 mo [55] studies (**Fig. 9B**). The lack of correlation between other studies and early developmental timepoints (1-2 mo [54]) suggests that hippocampal sex differences observed during postnatal development are distinct from those seen in adulthood and with aging.

**Fig. 9.**
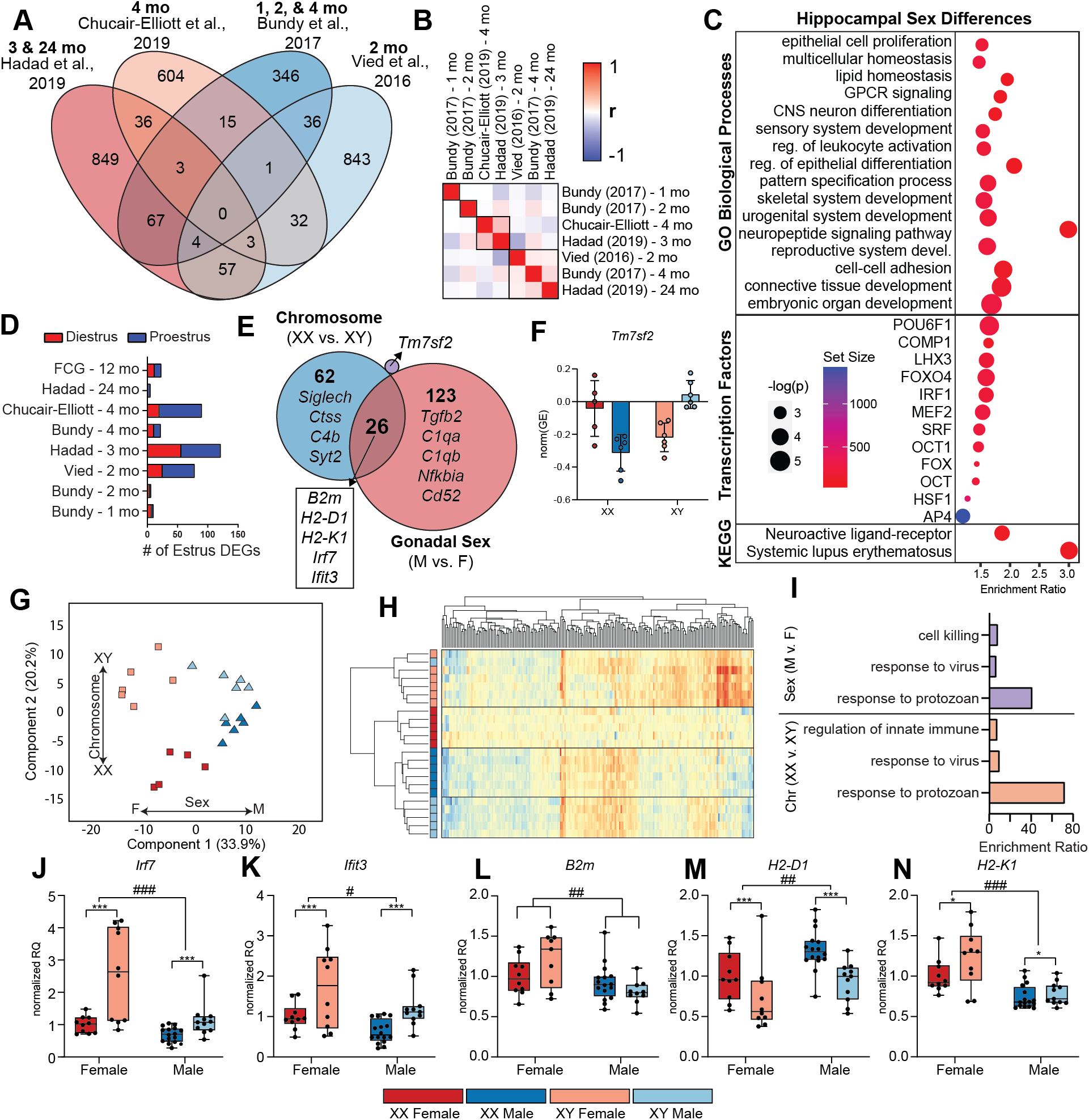
Transcriptomic analysis of autosomal differential expression from adult FCG hippocampi. DNA and RNA were isolated from FCG hippocampi (n=10-16/group). mRNA expression was assessed by stranded RNA-Seq (n=5-6/group) and RT-qPCR (n=10-16/group). Results were compared to previously published hippocampal transcriptomic sex differences. **Boxplots** represent median, interquartile range, and minimum/maximum normalized RQ. **A**) Comparison of four previous hippocampal transcriptomic studies (**Supplemental Table 4n**), shows 2896 autosomal-encoded sex differences in wild-type mice across all studies with no genes in common between all studies. **B**) Pairwise correlations of the fold-change (M v. F) for the 2896 autosomal DEGs were conducted for genes that were expressed in all studies. Hierarchical clustering of the correlation coefficients (Pearson r) shows grouping of studies. **C)** WebGestalt over-representation analysis of the 2896 autosomal-encoded sex differences in WT hippocampus identified KEGG pathways, transcription factors, and GO biological processes enriched in hippocampal sex difference genes (hypergeometric test, BHMTC, FDR<0.05). **D)** Previously identified estrus-cycle responsive genes (Jaric et al., 2019) were compared to hippocampal sex differences identified in previous WT (Fig. 9A) and the present FCG study. The number of estrus-responsive genes (diestrus female v. male – red, proestrus female v. male – blue) were plotted for each dataset. **E)** Differential expression analysis of FCG hippocampal RNA-Seq libraries identified 212 differentially expressed autosomal genes, 62 had a main effect of chromosome only (XX v. XY), 123 had a main effect of gonadal sex only (M v. F) and 26 genes had main effects of chromosome and gonadal sex. There was only one gene (*Tm7sf2*) which had an interactive effect of chromosomal and gonadal sex (Two-way ANOVA, BHMTC, FC>=1.25, FDR<0.1). **F)** Normalized gene expression of *Tm7sf2* by RNA-Seq (Two-way ANOVA, BHMTC, FC>=1.25, FDR<0.1, interactive effect). Bar chart is mean +/- SEM. **G)** Principal component analysis of differentially expressed autosomal genes showed separation of gonadal sex (M v. F) in the first component (33.9%) and separation of the chromosome (XX v. XY) in the second component (20.2%). **H)** Hierarchical clustering of differentially expressed autosomal genes shows separation of genotypes by gonadal sex and sex chromosome complement. **I)** ORA of the autosomal sex differences identified in the FCG hippocampus revealed biological differentially regulated by sex chromosome complement and/or gonadal sex. **J-N)** Differential expression of select genes was confirmed by RT-qPCR (n=10-16/group, Two-Way ANOVA, main effect of sex chromosome complement (XX v. XY) *p<0.05, **p<0.01, and ***p<0.001 or gonadal sex (M v. F) #p<0.05, ##p<0.01, and ###p<0.001). Only *Irf7* had a significant interaction between gonadal and chromosomal sex (p=0.012).

**Table 2.** Age, sample size, and estrus staging for previous hippocampal transcriptomic studies.

| Author/Year | Bundy et al., 2017 | Vied et al., 2016 | Chucair-Elliott et al., 2019 | Hadad et al., 2019 |
| --- | --- | --- | --- | --- |
| PMID | PMID: 28302071 | PMID: 26917114 | PMID: 31493147 | PMID: 31594536 |
| Data Availability | GEO Accession: GSE83931 | GEO Accession: GSE76567 | GEO Accession: GSE135752 | SRA bioProject: PRJNA523985 |
| Age (Group 1) | 1 mo | 2 mo | 4 mo | 3 mo |
| Age (Group 2) | 2 mo | - | - | 24 mo |
| Age (Group 3) | 4 mo | - | - | - |
| n | 5 | 5 | 3 | 6 |
| Estrus staging | 1 mo – no staging<br>2 mo – vaginal cytology; 2 estrus, 2 metestrus, 1 diestrus | No mention | No mention | No mention |

We next ran GO Biological Process, KEGG Pathway, and transcription factor over-representation analysis (ORA) using WEB-based GEne SeT AnaLysis Toolkit (WebGestalt, www.webgestalt.org) on autosomal genes that were differentially expressed by sex in at least one outside study (**Supplemental Table 2 (Union)**) (**Fig. 9C**, **Supplemental Table 6**, hypergeometric test, BHMTC, FDR<0.05). Enriched biological processes included lipid homeostasis, CNS neuron differentiation, regulation of leukocyte activation, pattern specification process, and reproductive system development, among others. Enriched transcription factors include immune regulators (IRF1, SRF, and MEF2), metabolic regulators (FOXO4), and vesicular trafficking (AP4), among others. Two significant KEGG pathways were over-represented in the hippocampal sex differences: neuroactive ligand-receptor interaction and systemic lupus erythematosus. Together, these data suggest that hippocampal sex differences alter neuronal differentiation/signaling and immune-related processes, which both could be partially mediated by differences in metabolic processes.

Of note is the fact that only one of the previous sex difference studies [54] mentioned estrus-cycle staging females, with collection of 2 estrus, 2 metestrus, and 1 diestrus females from each age group (1, 2, 4 mo). As such, this study [54] identified fewer estrus-dependent transcriptomic changes in 2 mo (6 genes) and 4 mo (22 genes) compared to age-matched 2 mo [57] (78 genes) and 4 mo [55] (90 genes) from other studies that did not estrus-cycle stage (**Fig. 9D, Table 2**). The 24 mo group [56] had few estrus-responsive DEGs, while the 2 mo [57], 3 mo [101], and 4 mo [55] that were not estrus cycle staged showed the largest number of estrus-responsive DEGs (**Fig. 9D**).

Next, we assessed autosomal sex differences in FCG hippocampus by sex chromosome complement (XX v. XY) and gonadal sex (M v. F) using directional RNA-Seq. After aligning, quantifying, and calling 212 differentially expressed genes: 62 genes differentially expressed by sex chromosome complement alone (XX v. XY), 123 genes differentially expressed by gonadal sex (M v. F), 26 genes differentially expressed by both sex chromosome complement and gonadal sex, and 1 gene with a significant interaction of gonadal and chromosomal sex (**Fig. 9E**). Principal component analysis of differentially expressed autosomal genes shows separation by gonadal sex in component 1 (33.9%) and separation by sex chromosome complement in component 2 (20.2%) (**Fig. 9G**). Hierarchical clustering of autosomal-encoded DEG shows proper clustering of samples by genotype (**Fig. 9H**). ORA of the 88 chromosomally-driven (XX v. XY) autosomal sex differences in the FCG hippocampus revealed four enriched biological processes: antigen processing and presentation, regulation of innate immune response, response to virus, and response to protozoan (Hypergeometric test, BH FDR<0.05). ORA of the 149 gonadal sex-driven (M v. F) autosomal sex differences in the FCG hippocampus revealed three enriched biological processes: leukocyte mediated immunity, response to virus, and response to protozoan (Hypergeometric test, BH FDR<0.05) (**Fig. 9I**).

Since we identified response to viruses and interferon-beta as pathways enriched in sex differences in the FCG hippocampus, we further examined interferon-associated genes IRF-7 and IFIT-3, both of which were differentially expressed by sex in other studies (**Supplemental Table 3 (Union)**). Interferon (IFN), as part of the innate immune system, triggers the expression of IFN-stimulated genes (ISGs) in response to viral recognition. IFN-beta is a type I IFN that is activated through PRRs [102], like *Tlr7*, is primarily in microglia [97]. In mouse models of AD, IFN was found to activate microglia leading to neuroinflammation and synaptic degradation. Blocking IFN signaling decreased microglia activation and concomitant synapse loss. Activation of IFN pathway was also observed in human AD [103]. Transcription factor IRF-7 is considered a “master regulator” in type-I IFN responses [104]. *Irf7* was differentially expressed in our study by sex chromosome complement (XX v. XY) and gonadal sex (M v. F) in FCG hippocampus as evidenced by RNA-Seq (**Supplemental Table 4**) and RT-qPCR confirmation (**Fig. 9J**). *Irf7* RT-qPCR also identified an interactive effect of gonadal and chromosomal sex (p=0.012). Interferon-induced protein with tetratricopeptide repeats 3 (IFIT3) is an antiviral RNA-binding protein which acts an intermediary in the activation of IRF-3 and upregulation of IFN-beta [105]. *Ifit3* was differentially expressed by sex chromosome complement (XX v. XY) and gonadal sex (M v. F) in FCG hippocampus as evidenced by RNA-Seq (**Supplemental Table 4**) and RT-qPCR confirmation (**Fig. 9K**).

Antigen processing and presentation pathways were also over-represented in our analysis of genes differentially expressed by chromosomal sex in the FCG hippocampus. Class I major histocompatibility (MHC-I) complexes function in innate viral antigen presentation and detection. Given that previous studies from our lab identified sexually divergent induction of MHC-I with aging in the mouse and rat hippocampus [106, 107], we examined expression of MHC-I components *B2m*, *H2-D1*, and *H2-K1*. Here we show differential expression of B2m by gonadal sex (**Fig. 9L**) and H2-D1/H2-K1 by sex chromosome complement and gonadal sex (**Fig. 9M-N**) in the FCG hippocampus (n=10-16/group, Two-Way ANOVA, main effect of sex chromosome complement (XX v. XY) *p<0.05, **p<0.01, ***p<0.001 and/or gonadal sex (M v. F) #p<0.05, ##p<0.01, ###p<0.001).

Based on the meta-analysis of previous studies, it appears that hippocampal autosomal sex differences are context-dependent and likely change throughout development and with aging. In the adult FCG hippocampus, sex differences were primarily enriched for in immune-related pathways and are regulated by both chromosomal and gonadal sex.

### Autosomal chromosome levels of methylation in FCG hippocampus by WGoxBS

After analyzing autosomal sex differences in FCG hippocampal gene expression, we assessed autosomal methylation in CG and CH context by WGoxBS. Overall, there were no differences in total autosomal mCG (**Fig. 10A**), as well as when split into in repetitive (**Fig. 10B**) and non-repetitive (**Fig. 10C**) element autosomal mCG. No difference was found in average autosomal mCH (**Fig. 10D**), repetitive element mCH (**Fig. 10E**) or non-repetitive element mCH (**Fig. 10F**). There were also no apparent differences in autosomal mCG patterning across CGI, shores, and shelves (**Fig. 10G**), gene bodies/flanking regions (**Fig. 10H**), or CTCF-binding sites/flanking regions (**Fig. 10I**). These data are consistent with previous findings that sex differences in mouse hippocampal autosomal methylation are site-specific and are a mix of both hyper- and hypomethylation in females compared to males [108].

**Fig. 10.**
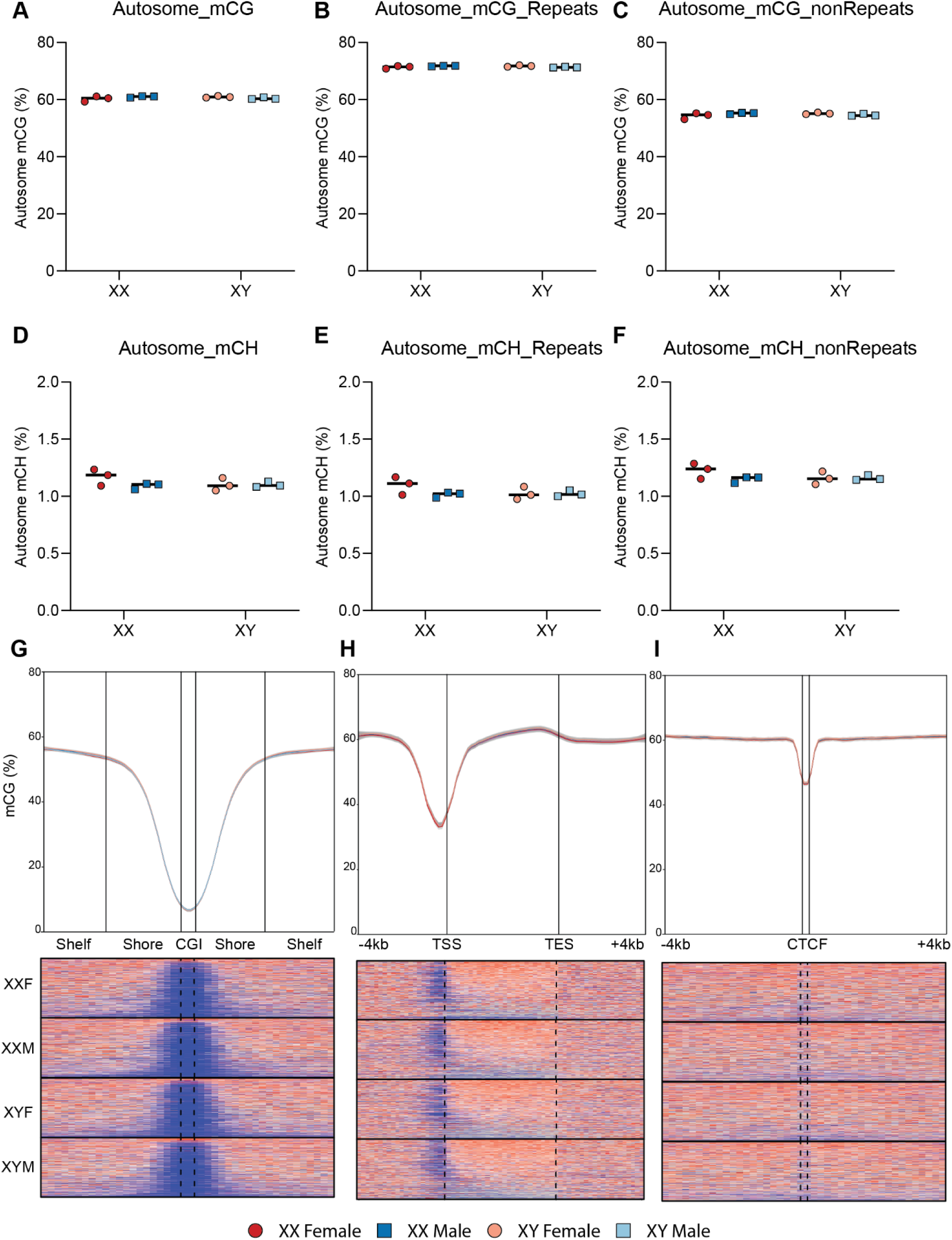
Autosomal levels of DNA methylation in FCG hippocampus by WGoxBS. DNA and RNA were isolated from FCG hippocampi (n=10-16/group). Autosomal methylation in CpG (CG) and non-CpG (CH) contexts were assessed by WGoxBS (n=3/group). A) There is no difference in overall mCG on autosomes, in either (B) Repetitive or (C) Non-repetitive elements. D) There is no difference in overall mCH on autosomes in either (E) Repetitive or (F) Non-repetitive elements (Two-way ANOVA). G-I) There is no difference in autosomal mCG in: (G) CpG Islands (CGI), shores, and shelves; (H) gene bodies and flanking regions (+/- 4 kb); or (I) CTCF binding sites and flanking regions (+/- 4 kb).

### Autosome differentially methylated regions (DMRs) by Whole Genome Oxidative Bisulfite Sequencing (WGoxBS) in FCG hippocampi

CG methylation was examined in the context of autosomal: CNS-associated histone marks [82] (H3K9ac, H3K4me2, H3K4me3, H3K27ac, H3K9me3, H3K36me3, H3K4me1, H3K27me3) (**Fig. 11A**), specific repetitive elements (SINE: Alu, B2, B4, MIR; LINE: L1, L2) (**Fig. 11B**), and proximal gene regulatory regions (enhancer, promoter, CGI, gene body) (**Fig. 11C**). There were no main effect differences by gonadal or chromosomal sex in average mCG across any of these genomic regions. In LINE elements (L1 & L2) there was a significant interactive effect of gonadal and chromosomal sex on autosomal LINE elements (L1, L2) mCG (Two-way ANOVA, p<0.05) but with a minimal methylation difference (<1%). While there were no sex effects in average mCG across these features, the levels of mCG varied among different features (i.e. Enhancers ∼ 48% and Promoters ∼3%).

**Fig. 11.**
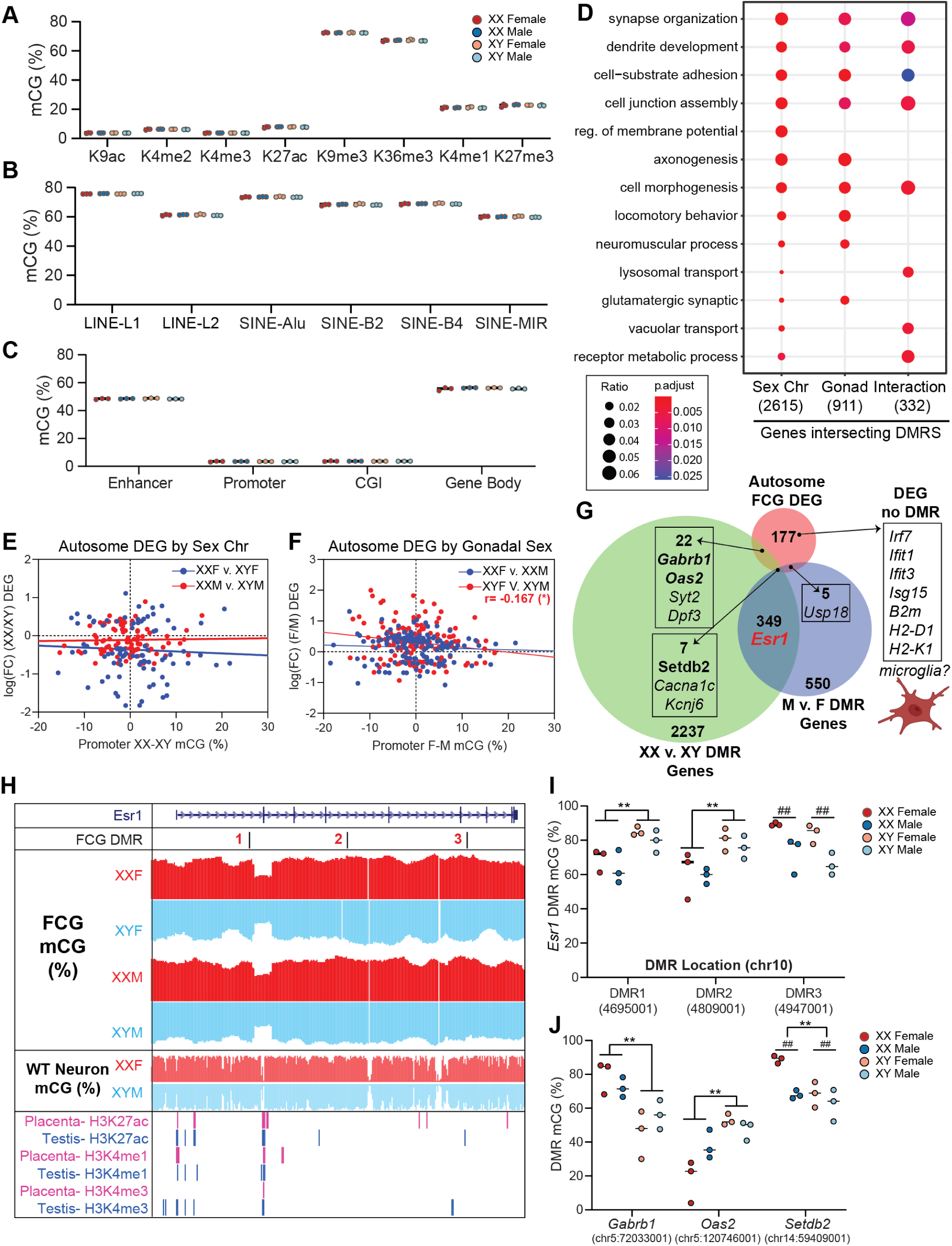
Epigenomic analyses of autosomal differentially methylated regions (DMRs) from adult FCG hippocampi. Autosomal methylation in CpG (CG) context was assessed by WGoxBS (n=3/group). First, average mCG levels in various genome contexts across autosomes were compared and no overall significant sex effects (gonadal or chromosomal) in mCG within: (**A**) autosomal P0 forebrain histone modification foci, (**B**) repetitive elements (LINE-L1, LINE-L2, SINE-Alu, SINE-B2, SINE-B4, SINE-MIR) or (**C**) autosomal proximal gene regulatory regions (Enhancer, promoter, CGI, Gene Body) (Two-way ANOVA). Differentially methylated regions (DMRs) (w=1000 bp) were called in methylKit (*post-hoc* Two-Way ANOVA with Bonferonni correction p<α=0.0083). **D)** GO Biological process over-representation analysis of autosomal genes (+/- 2kb) containing DMRs by sex chromosome, gonadal sex, or an interaction of gonadal and chromosomal sex. **E)** Correlation of promoter mCG (-2kb from TSS (XX-XY) and gene expression (log(FC)(XX/XY)) in autosomal DEGs by sex chromosome (XX v. XY) was not significant in XXF v. XYF or XXM v. XYM comparisons (Pearson r, p>0.05). **F)** Correlation of promoter mCG (-2kb from TSS (XX-XY)) and gene expression (log(FC)(XX/XY)) (Pearson r) in autosomal DEGs by sex chromosome (XX v. XY). Promoter methylation and gene expression were negatively correlated in XYF v. XYM comparison (Pearson r=-0.167, *p=0.045) but this was not observed in the XXF v. XXM comparison. **G)** Genes closest to autosomal DMRs (+/- 2kb gene body) with main effects of sex chromosome (XX v. XY) and/or gonadal sex (M v. F) were intersected with FCG autosomal DEGs (Fig. 9E). The majority (177/211=84%) of DEGs did not coincide with differentially methylated regions. **H)** Genome tracks of *Esr1* gene body (chr10) with FCG DMRs by chromosomal (1 & 2) and gonadal (3) sex, FCG mCG (%), WT male and female neuronal mCG (Lister et al., 2013), and male (testis) and female (placenta) specific histone peaks (H3K27ac, H3K4me1, H3K4me3) (The ENCODE Project Consortium et al. 2020). **I)** *Esr1* DMR (1, 2, & 3) mCG. DMRs 1 & 2 were differentially methylated by chromosomal sex (XX v. XY), while DMR 3 was differentially methylated by gonadal sex (M v. F) (Two-way ANOVA, main effect of sex chromosome **p<0.01, main effect gonadal sex, ##p<0.01). No significant interactions between gonadal and chromosomal sex were observed. **J)** DMR mCG levels within select autosomal FCG DEGs (*Gabrb1, Oas2*, *Setbd2*). DMRs within *Gabrb1* and *Oas2* were differentially methylated by sex chromosome complement (Two-way ANOVA, main effect sex chromosome complement, **p<0.01). The DMR within the *Setdb2* gene body was differentially methylated by chromosomal and gonadal sex (Two-way ANOVA, main effect sex chromosome complement, **p<0.01, main effect of gonadal sex, ##p<0.01). No interactive effect of gonadal and chromosomal sex was observed.

After exploring the overall levels and patterning of DNA methylation on the autosomes, autosomal mCG DMRs were assessed using 1 kb non-overlapping windows (Chisq-test, sliding linear model (SLIM) q<0.05) and *post-hoc* Two-way ANOVA with Bonferonni correction (p<α=0.0083) to assess main and interactive effects of chromosomal and gonadal sex. Using these criteria, we identified 4505 DMRs by sex chromosome (XX v. XY), 1385 DMRs by gonadal sex (M v. F), and 487 significant interactive effects of gonadal and chromosomal sex (**Supplemental Table 7**). We identified 2615 genic regions intersecting DMRs by chromosomal sex, 911 genic regions intersecting DMRs by gonadal sex, and 332 genic regions intersecting DMRs with an interactive effect of chromosomal and gonadal sex. These gene lists were assessed by GO Biological process over-representation analysis to identify significant pathways (**Supplemental Table 7**, hypergeometric test, BHMTC<0.2). ClusterProfiler [109] functional profiler assessment identified enrichment in neuronal-specific pathways (synapse organization, dendrite development, glutamatergic synaptic transmission) within the sex chromosomal and/or gonadal DMR-related genes (**Fig. 11D**).

To determine if there was a relationship between autosomal gene expression and promoter methylation, we correlated the difference in promoter methylation (mCG %) and differential gene expression (log(FC)) by sex chromosome (**Fig. 11E**, XX v. XY) and gonadal sex (**Fig. 11F**, M v. F). There was a weak negative correlation between differential gene expression and promoter methylation among the XYF v. XYM comparison (**Fig. 11F,** Pearson r=-0.167, *p=0.045). No other significant correlations were identified.

Most (84%) of the autosomal FCG DEGs from **Fig. 9E** did not overlap with a DMR (**Fig. 11G**). The autosomal DEGs that did overlap with DMRs appear to be primarily related to neuronal functions (*Gabrb1*, *Syt2*, *Cacna1c*, *Kcnj6*). The 177 DEGs not overlapping with DMRs, were enriched in immune-related genes (i.e. *Irf7*, *Ifit1/3*, *B2m*, *H2-D1*, *H2-K1*). Since immune cells, like microglia, are minority populations compared to neurons, the epigenomic signatures of microglia are not well represented in the bulk epigenomic data and require epigenomic analysis of these specific cell types to reveal methylation differences not observed in whole tissue analyses [110]. Further cell type-specific epigenomic data will need to be collected from the FCG model to determine the cell type-specific contributions to sex differential gene expression in the hippocampus.

Although *Esr1* was not differentially expressed in the FCG hippocampus, it was identified as a gene differentially expressed by sex in published mouse hippocampal data. Within the FCG mCG data, we identified 3 DMRs within the *Esr1* gene body DMRs by chromosomal (DMRs 1 & 2) and gonadal (DMR 3) sex (**Fig. 11H-I**). *Esr1* DMRs by chromosomal sex may suggest a sex differential imprinting mechanism used to regulate *Esr1* expression before the induction of gonadal hormones. The gonadal DMR indicates a potential feedback domain to modulate estrogen-receptor expression in response to gonadal hormones. Examples of differential methylation within differentially expressed genes was also shown for: GABA receptor *Gabrb1*, interferon response gene *Oas2*, and methyl-CG dependent histone modifier *Setdb2* (**Fig. 11J)** demonstrating that while some DEGs do not have a correspond methylation difference, in some genes like these there is a corresponding methylation change.

Despite a strong immune-related transcriptomic signature, differential methylation appears to be mostly involved in neuron-related pathways. This could be expected given that neurons are the predominant cell type in the hippocampus. And while there is variable association of DMRs to DEGs, the expression analysis is examining steady state gene expression and these DMRs may cause differential gene expression in acute, stimulus-responsive situations. Future studies will also need to specifically examine minority microglia cell population to uncover methylation-gene expression relationships in these cells.

## Discussion

The study of sex effects in brain health and disease have begun receiving needed experimental attention in neuroscience studies. Not only do the sexual dimorphisms, differences, and divergences [111] need to be characterized but also the regulatory mechanisms giving rise to these sex effects. While hormonal mechanisms (both organizational and activational) have been the most studied, the potential regulation of sex effects by sex chromosomes, either independently or in concert with gonadal hormones, has received relatively limited attention. Symptomology associated with human sex chromosome aneuploidies [39–41] and experimental evidence obtained from FCG mouse studies [17, 31, 32, 48, 53] have provided critical insight into the importance of sex chromosomes (independent of gonadal sex) to sex effects seen in development, aging, and disease. Previous reports have established sex differences in the hippocampal epigenome and transcriptome with brain aging [108, 125] and disease [57, 126, 127], further suggesting that the intersection of epigenetic alterations and sex effects requires further investigation. Here we use the FCG mouse model to delineate the contributions of gonadal and chromosomal sex to the epigenomic regulation of hippocampal gene expression in adult mice.

We first validated the genetics of the FCG model, confirming prior imaging and PCR studies [53] and providing a precise location of *Sry* insertion to an intergenic region of Chr3. While there are ∼13 copies of *Sry*, the insertion is not within an annotated gene. Since the concatemer of *Sry* could disrupt the chromatin accessibility within and around the insertion site on chromosome 3, we assayed for ectopic *Sry* expression and changes to gene expression genes adjacent to the insertion site. However, we found no evidence of *Sry* expression in the hippocampus (as expected) and no disruption of gene expression in adjacent genes. Additionally, tissue-specific expression of Y-encoded genes (*Kdm5d*, *Eif2s3y*, *Ddx3y*, *Uty*) was maintained in the FCG, despite deletion of *Sry*. Together these data provide confidence that the transgenic manipulations used to generate the FCG are not confounding the analysis of sex differences in the hippocampus.

XCI, the random silencing of one X-chromosome (Xi) during female embryogenesis, is an important mechanism for gene dosage compensation between males and females [112]. Maintenance of XCI occurs via intricate epigenomic mechanisms, including DNA hypermethylation, lncRNA expression (i.e. *Xist*), and specific histone modifications [113]. However, Xi is not fully repressed and a subset of X-chromosome genes consistently escape XCI, leading to higher expression in females [114, 115]. Escape from X-inactivation in females has been implicated in the female bias in autoimmune diseases [116, 117] and AD [118]. X-chromosomal paired transcriptomic and DNA methylation sequencing revealed that sex effects in XCI and escape are principally driven by sex chromosome content, but may be “fine-tuned” by gonadal sex. Contributions of gonadal sex XCI escape may explain why escape genes vary by tissue [58]. We found that methylation of ChrX proximal gene regulatory features (gene body, promoter, enhancer, CGI) was regulated autonomously by the sex chromosomes with patterning consistent with XCI signatures previously reported in WT mice [55, 77]. Among ChrX DEGs, promoter methylation showed a strong negative correlation to gene expression, suggesting an important role of DNA methylation in modulating escape from XCI. We also propose a chromatin signature for genes that always escape XCI, which is distinct from the chromatin context of variable escape genes. This led us to hypothesize that variable XCI escape may result from sex hormonal dependent mechanisms.

Repetitive elements (especially LINE-L1) have been implicated in XCI spreading, escape, and maintenance [79, 119–121]. Specific L1 element expression is posited to facilitate propagation of XCI [119]. In the FCG hippocampus, L1 elements were the only X-chromosomal genomic features that showed lower mCG in XX (v. XY) genotypes. The hypomethylation of XX (v. XY) ChrX L1 elements could represent a sex chromosome autonomous mechanism of XCI. The only ChrX genomic feature that displayed differential mCG by gonadal sex was SINE-Alu repeats, with higher methylation in gonadal females than males. Using *Aff2* as an example, we propose an Alu-repeat dependent mechanism for epigenomic modulation of gene expression by gonadal sex. The idea of hormonal tuning of XCI is bolstered by ChrX chromatin accessibility alterations by estrus-cycle stage [90]. Further hormone manipulation studies (i.e. ovariectomies) will need to be conducted to confirm this hypothesis.

Autosome-encoded hippocampal gene expression and methylation sex differences are regulated by both sex chromosomes and gonadal sex. Importantly, hippocampal changes induced with age [56, 106–108] and female estrus-cycle stage [90, 122, 123] need to be considered in calling consistent gene expression differences driven by gonadal sex. Despite the FCG chromosomal and gonadal sex differential gene expression being strongly enriched in inflammatory pathways (i.e. MHC-I, interferon-response), regions of differential methylation were strongly associated with neuronal functions. Additionally, comparison of gene expression and promoter methylation within DEGs did not show the strong, negative correlations normally seen with methylation-dependent gene transcription. Since microglia are a minority cell population in the hippocampus, changes in microglial methylation patterns are difficult to detect in bulk sequencing data. Thus, further studies examining cell type-specific epigenomic regulation of sex differential gene expression are needed.

The mechanism for sex chromosome content directly regulating the autosomes may lie, in part, in the X-chromosome encoded histone demethylases *Kdm6a* and *Kdm5c* and Y-encoded *Uty* and *Kdm5d* since sex chromosome-regulated DEGs were enriched by H3-K27 and H3-K4 demethylation peaks. Recent reports have highlighted the importance of sex chromosome-encoded histone modifiers in mouse models of autoimmunity [124, 125], adiposity [126], and AD [32]. Further work will be needed to manipulate individual X- and Y- encoded histone modifiers to determine their effect on the epigenomic landscape of both sex chromosomes and autosomes in the context of brain aging and disease.

Of particular relevance to the present hippocampal study, Davis et al. (2020) used the FCG model to establish that a second X-chromosome confers protection against AD [32] through X-linked H3-K27 demethylase *Kdm6a*. They showed that hippocampal overexpression of *Kdm6a* in male mice with a familial AD mutation (XY-hAPP-J20) attenuated AD-induced cognitive deficits when compared to controls. This study provides critical insight into the sex-biased regulation of AD pathogenesis and opens several new lines of investigation. Primarily, do alterations in H3-K27 methylation play a causal role in the neuroprotective effects of *Kdm6a*? If so, are alterations in H3-K27 methylation localized to the X-chromosome or do sex chromosome-encoded histone modifiers alter the autosomal epigenomic landscape? Do other cell types (e.g., microglia) rely on *Kdm6a* for proper function in response to AD pathology? In the hAPP-J20 model used [30], the humanized APP is inserted into a noncoding portion of Zinc finger protein *Zbtb20* gene (involved in interferon response in macrophages [55]), causing decreased hippocampal *Zbtb20* expression [56]. Since interferon response genes (e.g., *Tlr7, Irf7, Ifit3*) were identified in the present study as being differentially regulated by both sex chromosome complement and gonadal sex, a different mouse model of AD (in lieu of the hAPP-J20) may allow for further identification of sex differentially regulated microglial inflammatory pathways contributing to AD.

Taken together these findings are consistent with the hypothesis that the sex chromosomes have effects on the hippocampal epigenome and transcriptome independent of gonadal sex. This is the first demonstration of the separable effects of hormones and sex chromosomes on the epigenome and transcriptome in the FCG hippocampus. However, a number of questions remain to be answered in future studies. These principally consist of further controlling for gonadal hormone status but analyzing FCG mice that have been gonadectomized after development at ∼1-2 months of age. This will control for any activational hormonal differences between the genotypes. It is worth noting that these studies were conducted in adult mice ∼12 months of age.

Most importantly, analyses of specific cell types are needed for future studies, as transcriptomic and epigenomic profiles differ between neuronal, glial, and other cell types of the CNS. Expanding the research on the epigenetic and transcriptomic landscapes in specific cell populations of the brain will help us more clearly delineate cellular contributions to sex differences in future molecular studies. Further investigation is highly warranted given the significant effects of sex chromosome regulation of gene expression and DNA modification patterns in cis of the X chromosome and in trans of the autosomes.

## Methods

### Animals

All animal procedures were approved by the Institutional Animal Care and Use Committee at the University of Alabama at Birmingham (UAB) under protocol 21506. Four Core Genotypes mice on a C57Bl/6J background were obtained from the Jackson Laboratory (Stock #010905) where they were revitalized from frozen embryo stocks. Breeder pairs were set up with a XX female and a XY male, as described in the introduction. Pups were weaned into sex specific cages of 4-5 animals based on visual inspection of genitalia. Animals were maintained in an AAALAC approved UAB animal facility at 21°C on a 12/12 light/dark cycle (lights on at 6:00am). Animals were provided ad libitum water and standard mouse chow (NIH31) until time of tissue collection, ∼12 months of age for both males and females. Euthanasia prior to tissue harvesting was carried out by cervical dislocation followed by rapid decapitation. Genotyping was performed using the primers: Sry (5′-AGC CCT ACA GCC ACA TGA TA-3′, 5′-GTC TTG CCT GTA TGT GAT GG-3′), Ymt (Y chromosome-specific sequence, 5′-CTG GAG CTC TAC AGT GAT GA-3′, 5′-CAG TTA CCA ATC AAC ACA TCA C-3′), and myogenin (5′-TTA CGT CCA TCG TGG ACA GCA T-3′, 5′-TGG GCT GGG TGT TAG TCT TAT-3′) [53], following Jackson Laboratory’s standard PCR assay genotyping protocol #5590.

### High molecular weight (HWM) DNA isolation for pseudo long-read genomic sequencing

Hippocampi were dissected from a FCG XYM mouse, snap frozen in microfuge tubes with liquid nitrogen, and stored at -80°C prior to DNA isolation. HMW DNA was extracted from fresh-frozen tissue according to 10X Genomics sample preparation protocol (https://support.10xgenomics.com/genome-exome/sample-prep/doc/demonstrated-protocol-hmw-dna-extraction-from-fresh-frozen-tissue). Frozen tissue was thawed on ice and minced with a razor blade. The tissue was gently Dounce homogenized in 500 µL of Nuclei Isolation Buffer (Sigma Nuclei PURE Prep Kit: Lysis Buffer, 1 mM DTT, Sigma Nuclei PURE Prep Kit: 10% Triton X-100). After a brief centrifugation, the supernatant was transferred using a wide-bore pipette tip to a 2.0 mL round-bottom tube and centrifuged at 500 x g for 5 minutes. The supernatant was discarded, and pelleted nuclei were then resuspended in 70 µL of ice-cold PBS. To digest the nuclei, 10 µL of Proteinase K was added to the resuspended nuclei, followed by 70 µL ice-cold Digestion Buffer (20 mM EDTA, pH 11, 2mM Tris-HCl, pH 8.3,10 mM N-Laurylsarcosine sodium salt). Samples were rotated end-over-end for 2 hours at room temperature. To purify the DNA, Tween-20 was added to the sample to a final concentration of 0.1% and then 1X SPRISelect Reagent was added. The samples were rotated end-over end for 20 min. Tubes were placed in the DynaMag-2 magnetic rack to allow bead capture. After removing and discarding the supernatant, the beads were washed twice with 70% ethanol. The DNA was eluted from the beads with 50 µL Sample Elution Buffer (Qiagen AE Buffer, 0.1% Tween-20). Qubit dsDNA BR kit was used to quantify the DNA.

### 10X Linked-Read Library Preparation

Linked-read genomic libraries Chromium were constructed from 1 ng of HMW DNA from an XYM using Chromium Genome Library Prep Kit (#PN-120229, 10X Genomics, Pleasanton, CA), according to manufacturer’s instructions. Briefly, following HMW gDNA extraction 1 ng of HMW DNA was loaded onto a Chromium Genome Chip Kit (#PN-120216, 10X Genomics) for Gel Bead-in-Emulsions (GEM) generation and barcoding. After SPRISelect bead cleanup and library construction, libraries were normalized to 4 nM, pooled, and sequenced NextSeq500 (High PE150) in the OMRF Clinical Genomics Center (CGC). Sequencing data was aligned to the mm10 genome using the LongRanger pipeline (10X Genomics) and visualized in the Loupe browser.

### Isolation of DNA/RNA from FCG hippocampi

Hippocampi were dissected from XXF, XXM, XYF, and XYM FCG mice (n=10-16/group), snap frozen in microfuge tubes with liquid nitrogen, and stored at -80°C prior to DNA isolation. Nucleic acids (DNA/RNA) were isolated from flash-frozen tissues using the All Prep DNA/RNA Mini Kit (Qiagen, Germantown, MD) as previously described [127–129]. Briefly, 600 µL of Buffer RLT with beta-mercaptoethanol was added to the tube containing the frozen hippocampi. A steel bead was added to the tube and homogenized for 30 s at 30 Hz using a Tissue Lyser II (Qiagen). Homogenate was loaded onto a DNA spin column and the flow through supplemented with ethanol was loaded onto an RNA spin column. Columns were washed and nucleic acids eluted by centrifugation. Total DNA/RNA were quantified by Nanodrop (Thermofisher Scientific, Madison, USA). Quality of DNA and RNA were assessed by genomic and RNA screentapes, respectively, on a Tapestation 2200 (Agilent Technologies, Frankfurt, Germany). Only samples with RNA and DNA integrity numbers > 7 were used for subsequent experiments.

### Digital PCR (dPCR) *Sry* copy number assay

Custom *Sry* fluorogenic copy number assays were designed and purchased from Integrated DNA Technologies (Coralville, IA) with primers (5’-GTC ACA GAG GAG TGG CAT TT-3’, 5’-AGT CTT GCC TGT ATG TGA TGG-3’) and probe (5’-/56-FAM/CAACCTTCT/ZEN/GCAGTGGGACAG GAA/3IABkFQ/-3’). Mouse *Tert* fluorogenic copy number assay was used as a single copy gDNA control as reference (#4403316, Life Technologies). DNA from FCG hippocampi (n=3/group) was used for dPCR, as previously [129], using the QuantStudio 3D Digital PCR kit (#A26361, ThermoFisher Scientific), according to manufacturer’s instructions. After combining DNA with the Quantstudio 3D mastermix and *Sry* or *Tert* fluorogenic assay, reactions were loaded onto a Quantstudio 3D PCR chip with a Quantstudio 3D chip loader (#4482592, Thermofisher Scientific) and cycled on a GeneAmp PCR system 9700 with a flatblock attachment. Chips were read by the Quantstudio 3D chip reader (#4489084, Thermofisher Scientific) and analyzed using Quantstudio 3D AnalysisSuite cloud software V3.1.

### Quantitative PCR (qPCR)

Confirmation of gene expression levels was performed with qPCR as previously described [55, 130, 131]. cDNA was synthesized with the ABI High-Capacity cDNA Reverse Transcription Kit (Applied Biosystems Inc., Foster City, CA) from 25ng of purified RNA. qPCR was performed with gene-specific primer probe fluorogenic exonuclease assays (TaqMan, Life Technologies, Waltham, MA, **Supplemental table 5**) and the QuantStudio 12K Flex Real-Time PCR System (Applied Biosystems). Relative gene expression (RQ) was calculated with Expression Suite v 1.0.3 software using the 2^−ΔΔ^Ct analysis method with GAPDH as an endogenous control. Statistical analysis of the qPCR data was performed using GraphPad Prism 8 (San Diego, CA). Two-way ANOVA analyses were performed followed by the Tukey’s multiple comparison test (p<0.05).

### Library construction and RNA sequencing (RNA-seq)

Illumina’s TruSeq Stranded mRNA Library Prep Kit (#20020594, Illumina) was used on 500 ng of total RNA for the preparation of strand-specific sequencing libraries according to manufacturer’s guidelines. As previously described [56], rRNA depletion was performed prior to library construction.

RNA was isolated from fresh-frozen hippocampal tissue of 12 mo FCG mice (n=5-6/group), using Qiagen AllPrep DNA/RNA Mini Kit. After verifying RNA integrity numbers (RIN) with TapeStation (Agilent) and quantifying RNA with Qubit dsDNA Broad Range Assay kit (Invitrogen), 1 ug of RNA was used to construct RNA-Seq libraries using the Illumina TruSeq Stranded RNA Library Prep Kit, following the manufacturer’s guidelines. cDNA libraries were sized by TapeStation (Agilent) and quantified by qPCR (KAPA Biosystems). Libraries were then normalized to 4 nM, pooled, denatured, and diluted for sequencing on Illumina Hiseq2500 in a 2x100 bp fashion.

### RNA-Seq Data Analysis

Following sequencing, reads were trimmed, aligned, differential expression statistics and correlation analyses were performed in Strand NGS software package (Agilent), as previously described [55]. Reads were aligned against the Mm10 build of the mouse genome (2014.11.26). Alignment and filtering criteria included: adapter trimming, fixed 2bp trim from 5’ and 6bp from 3’ ends, a maximum number of one novel splice allowed per read, a minimum of 90% identity with the reference sequence, a maximum of 5% gap, trimming of 3’ end with Q<30. Alignment was performed directionally with Read 1 aligned in reverse and Read 2 in forward orientation. Reads were filtered based on the mapping status and only those reads that aligned normally (in the appropriate direction) were retained. Normalization was performed with the DESeq algorithm [132]. Transcripts with an average read count value >20 in at least 100% of the samples in at least one group were considered expressed at a level sufficient for quantitation per tissue. Those transcripts below this level were considered not detected/not expressed and excluded, as these low levels of reads are close to background and are highly variable. A fold change >|1.25| cutoff was used to eliminate those genes which were unlikely to be biologically significant and orthogonally confirmable due to their very small magnitude of change. For statistical analysis of differential expression, a two-way ANOVA with the factors of sex chromosome complement and gonadal sex and a Benjamini-Hochberg Multiple Testing Correction (BHMTC) with an FDR of 0.1. Visualizations of hierarchical clustering and principal components analysis were performed in Strand Next Generation Analysis Software (NGS) (Version 3.1, Bangalore, India). The entirety of the sequencing data is available for download in FASTQ format from NCBI Gene Expression Omnibus (GEO) under accession number GSE184098.

### Bisulfite amplicon sequencing (BSAS)

BSAS for specific gene promoters was performed as previously described [110, 128]. Briefly, FCG hippocampal DNA (200 ng) was bisulfite converted with the EZ DNA Methylation-Lightning Kit (#D5030T; Zymo Research, Irvine, CA), according to the manufacturer’s guidelines. Bisulfite-PCR was conducted using primer sets (Integrated DNA Technologies; **Table 1**) designed against specific genomic regions with Methyl Primer Express v1.0 software (Thermofisher Scientific). PCR amplicons were cleaned with AmpureXP beads (#A63882; Beckman Coulter Life Sciences, Indianapolis, IN) using size selection with 0.7x bead ratio. Following clean-up, the amplicons were quantified using QubitTM dsDNA HS assay kit (#Q32851; Thermofisher Scientific) and pooled. One ng of the pooled amplicons was used for library construction with the Nextera XT DNA library preparation kit (#FC-131–1096; Illumina, San Diego, CA), according to the manufacturer’s instructions. Libraries were quantified with Qubit dsDNA HS assay kit and TapeStation HD1000, normalized to 1 nM or 4 nM, and pooled for sequencing. Pooled libraries were then sequenced on iSeq or MiSeq (Illumina) at loading concentrations 35 pM or 8 pM, respectively. Fastq files were aligned to amplicon sequences in CLC Genomics Workbench 11.0 (Qiagen) using the “Map Bisulfite Reads to Reference” feature. Site-specific CpG (CG) and CH methylation percentages were extracted for downstream analysis.

### Library construction and oxidative bisulfite sequencing (WGoxBS-seq)

DNA was isolated from fresh-frozen hippocampal tissue of 12 mo FCG mice (n=3/group), using Qiagen AllPrep DNA/RNA Mini Kit. Whole genome oxidative bisulfite sequencing libraries were prepared according to manufacturer’s guidelines (Ovation Ultralow Methyl-Seq Library System, Tecan Genomics, Inc., Redwood City, CA) as previously described [55, 110]. Briefly, 1 µg of gDNA in 50 µl 1X low-EDTA TE buffer was sheared with a Covaris E220 sonicator (Covaris, Inc., Woburn, MA) to an average of 200 base pairs. Sheared products were sized by capillary electrophoresis (DNA HSD1000, Agilent) and cleaned using an Agencourt bead-based purification protocol. After quantifying (Qubit dsDNA, Thermofisher Scientific) the cleaned DNA, 200 ng gDNA fragments were prepared in a 12 µl volume to which 1µl of spike-in control DNA (0.08 ng/ul) with known levels of specific mC, hmC, and fC at individual sites was added. End repair, ligation of methylated adaptors (#L2V11DR-BC 1-96 adaptor plate, Tecan Genomics) and final repair were performed according to manufacturer’s instructions. Normalized DNA was oxidized and then bisulfite-converted with the True Methyl oxBS module (NuGEN) with desulfonation and purification. 22 µl of libraries were eluted from the magnetic beads. qPCR was used to determine the number (N) of PCR cycles required for library amplification. Oxidative bisulfite-converted samples were amplified for 13 cycles [95° C- 2 min, N (95°C-15 s, 60°C-1 min, 72° C-30s)]. Amplified libraries were purified with Agencourt beads and eluted in low-EDTA TE buffer. Capillary electrophoresis (TapeStation HSD1000, Agilent) was used to validate and quantify libraries. Amplified libraries were normalized to a concentration of 4 nM and pooled, denatured, and diluted to 12 pM for sequencing on NovaSeq 6000 (Illumina) according to manufacturer’s guidelines with the exception of a custom sequencing primer (MetSeq Primer) that was spiked in with the Illumina Read 1 primer to a final concentration of 0.5 µM.

### OxBS-seq data analysis

Global levels of mCG, hmCG, and mCH were analyzed as previously described [55],[110] Before aligning, paired-end reads were adaptor-trimmed and filtered using Trimmomatic [133] 0.35. End-trimming removed leading and trailing bases with Q-score<25, cropped 4 bases from the start of the read, dropped reads less than 25 bases long, and dropped reads with average Q-score<25. Unpaired reads after trimming were not considered for alignment. Alignment of trimmed OxBS-converted sequences was carried out using Bismark [134] 0.16.3 with Bowtie 2 [135] against the mouse reference genome (GRCm38/mm10). Bams were de-duplicated using Bismark. Methylation call percentages for each CpG and non-CpG (CH) site within the genome were calculated by dividing the methylated counts over the total counts for that site in the oxidative bisulfite - converted libraries (OXBS). Genome-wide CpG and CH methylation levels were calculated separately. BAM files generated during alignment were run through methylKit in R [136] to generate context-specific (CpG/CH) coverage text files. Bisulfite conversion efficiency for C, mC, and hmC was estimated using CEGX spike-in control sequences. Untrimmed fastq files were run through CEGX QC v0.2, which output a fastqc_data.txt file containing the conversion mean for C, mC, and hmC. The entirety of the sequencing data is available for download in FASTQ format from NCBI Gene Expression Omnibus (GEO) under accession number GSE184013.

### DMR Analysis

CpG text files were read into methylKit [136] and converted to an object. The mouse genome was tiled in 1000 nt non-overlapping windows. Each window was filtered for a minimum count of 10. Samples were then united and compared for windows covered in at least two samples per group. Differentially methylated regions (DMRs) were called using default parameters. DMRs were filtered to differences that were >10% different between at least two groups and had a SLIM-generated q-value less than 0.05. The methylDiff object was intersected with the methylBase object to calculate the % methylation for each window that passed the described filtering. *Post-hoc* assessment of main and interactive effects of chromosomal and gonadal sex was conducted using a Two-way ANOVA with Bonferonni correction for six pairwise comparisons (p<α=0.0083).

### Software usage for tertiary analysis of transcriptomic and epigenomic data

Over-representation analysis (ORA) for GO Biological processes, transcription factor networks, and KEGG pathways was conducted using WEB-based GEne SeT AnaLysis Toolkit (WebGestalt, www.webgestalt.org) [137]. EnrichedHeatmap [138] in R was used to visualize DNA methylation levels across ChrX and autosomal CGI, Gene body/promoter, and CTCF binding sites. Bed files for genomic features were downloaded from the UCSC table browser [139] and visualized with custom tracks using the UCSC genome browser (http://genome.ucsc.edu) [140]. BEDtools [141] was used to intersect mCG/CH text files with specific genomic features. TrawlerWeb [142] was used to identify over-represented motifs from BED files. STRING protein-protein association networks [81] were used to determine potential protein interactors with SP1. Venny [143] and BioVenn [144] were used to compare gene lists and make Venn diagrams. BedSect [145] was used to create Jaccard distance matrices comparing genomic features to differentially methylated and expressed regions. ggplot2 [146] and clusterProfiler [147] were used to generate dotplots of GO biological processes.

## Acknowledgments

This work was supported by grants from the National Institutes of Health (NIH) P30AG050911, R01AG059430, R56AG067754, T32AG052363, F31AG064861, P30EY021725, P30AG050886, R21AG058811, R01AG057434, R01AG070035, R01AG069742, Oklahoma Center for Adult Stem Cell Research (OCASCR), a program of the Oklahoma Tobacco Settlement Endowment Trust, BrightFocus Foundation (M2020207), and Presbyterian Health Foundation. This work was also supported in part by the MERIT award I01BX003906 and a Shared Equipment Evaluation Program (ShEEP) award ISIBX004797 from the United States (U.S.) Department of Veterans Affairs, Biomedical Laboratory Research and Development Service. The authors would also like to thank the Clinical Genomics Center (OMRF) for assistance and instrument usage. The authors also acknowledge Robyn Berent (administrative support and lab management), Adeline Machalinski (animal colony management), Ashley Martin (manuscript review), and Hunter Porter (scientific discussions).

## Declarations

### Availability of data and material

The datasets generated during and/or analyzed during the current study are available in the NCBI Gene Expression Omnibus (GEO) repository and Sequence Read Archive (SRA) with accession numbers: GSE83931, GSE135752, GSE76567, GSE184098, and PRJNA523985. All other data are available from the corresponding author on reasonable request

### Code availability

All data were analyzed with commercially available software packages and open-source web applications and R packages, as indicated in the text.

### Author contributions

Sarah R. Ocañas: first author, design of the study, execution of experiments, data acquisition, analysis, and interpretation, figure generation, manuscript writing and preparation.

Victor A. Ansere: execution of experiments, data acquisition, analysis, and interpretation, figure generation, manuscript preparation.

Kyla B. Tooley: execution of experiments, data acquisition, analysis, and interpretation, figure generation.

Niran Hadad: data analysis and interpretation, figure generation, manuscript writing and preparation.

Ana J. Chucair-Elliott: design of the study, data interpretation, manuscript writing and preparation.

David R. Stanford: design of the study, data analysis and interpretation.

Shannon Rice: execution of experiments, data acquisition, analysis, and interpretation.

Benjamin Wronowski: execution of experiments, data acquisition, analysis, and interpretation, figure generation.

Kevin D. Pham: data analysis and interpretation, manuscript writing and preparation.

Jessica M. Hoffman: design of the study, execution of experiments, data analysis and interpretation, manuscript writing and preparation.

Steven N. Austad: design of the study, data analysis and interpretation, manuscript writing and preparation.

Michael B. Stout: design of the study, execution of experiments, data acquisition, analysis, and interpretation, manuscript writing and preparation.

Willard M. Freeman: Corresponding author, design of the study, data analysis and interpretation, figure generation, manuscript writing, preparation, and submission.

### Ethics approval

Not applicable

### Consent to participate

Not applicable

### Consent for publication

Not applicable

